# Cellular Mechanisms Underlying Central Sensitization in a Mouse Model of Chronic Muscle Pain

**DOI:** 10.1101/2022.03.22.485320

**Authors:** Yu-Ling Lin, Zhu-Sen Yang, Wai-Yi Wong, Shih-Che Lin, Shuu-Jiun Wang, Shih-Pin Chen, Jen-Kun Cheng, Hui Lu, Cheng-Chang Lien

## Abstract

Chronic pain disorders are often associated with psychiatric symptoms. The central nucleus of the amygdala (CeA) has emerged as an integrative hub for nociceptive and affective components during the development of central pain. Although the exact cause for this process remains unknown, prior adverse injuries are precipitating factors and thought to transform nociceptors into a primed state for chronic pain. However, the cellular basis underlying the primed state and the subsequent pain chronification remains unknown. Here, we investigated cellular and synaptic alterations of the CeA in a mouse model of chronic muscle pain. In these mice, local infusion of pregabalin, a clinically approved drug for fibromyalgia and other chronic pain disorders, into the CeA or selective inactivation of somatostatin-expressing CeA (CeA-SST) neurons during the priming phase prevented pain chronification. Further, electrophysiological recording revealed that CeA-SST neurons received increased excitatory transmission and showed enhanced excitability in chronic pain states. In line with the possible role of CeA-SST neurons in central sensitization, chemogenetic inactivation of CeA-SST neurons or pharmacological suppression of nociceptive afferents from the brainstem to CeA-SST neurons by pregabalin after the development of chronic muscle pain alleviated pain and negative emotions.

## Introduction

Prior adverse events or perceived physical injuries are considered to be one of precipitating factors for chronic pain (Althaus et al., 2012; M. Aronoff and B. Feldman, 2009; Mills et al., 2019; Stevans et al., 2021). A prior injury is thought to transform nociceptors into a primed state at the cellular level, which may last variably in different types of chronic pain disorders (Kandasamy and Price, 2015; Reichling and Levine, 2009; Sun and Chen, 2016). Subsequent injuries occurring during the priming phase can lead to pain chronification (Sun and Chen, 2016). Acid-induced muscle pain (AIMP) in rodents, which is considered as a preclinical model for chronic muscle pain (MP) disorders, including fibromyalgia syndrome, requires two episodes of acute pain induction (Sluka et al., 2001). Specifically, the second acid injection into the gastrocnemius muscle of animals during a priming phase is required for the development of chronic muscle pain (Chen et al., 2014; Sluka and Clauw, 2016; Sun and Chen, 2016). Chronic pain disorders are associated with psychiatric comorbidities (Mills et al., 2019; Romano and Turner, 1985). Indeed, patients with fibromyalgia syndrome not only suffer from chronic widespread pain, but are frequently associated with comorbid psychological, sleep, and cognitive disturbances (Clauw, 2014; Sluka and Clauw, 2016).

Central sensitization is believed to underlie widespread MP and affective symptoms in fibromyalgia syndrome (Fu et al., 2008; Pedersen et al., 2007; Sheng et al., 2017; Woolf, 2011). However, the cellular basis underpinning central sensitization in chronic MP disorders is poorly known. Among the anatomically and functionally distinct amygdaloid nuclei (Duvarci and Pare, 2014; LeDoux, 2000), the central nucleus of the amygdala (CeA) has emerged as an integrative hub for nociceptive and affective components during the development of chronic pain states (Neugebauer et al., 2004; Thompson and Neugebauer, 2017). The CeA receives a direct nociceptive projection from the parabrachial nucleus (PBN). Maladaptive changes in synaptic transmission at the PBN to CeA neuron synapses are thought to underlie persistent pathological pain states (Ikeda et al., 2007; Li and Sheets, 2020; Wilson et al., 2019). The CeA houses distinct GABAergic neuronal populations and the two major groups are the somatostatin-expressing (SST) and the protein kinase C delta-expressing (PKCδ) neurons. These two types of neurons make reciprocal inhibition. It is generally believed that PKCδ neurons are pronociceptive whereas SST neurons are antinociceptive under physiological conditions (Wilson et al., 2019). However, the role of these two types of neurons involved in the regulation of nociception are inconsistent in different chronic pain models (Wilson et al., 2019; Zhou et al., 2019).

In this study, we aimed to investigate the cellular basis underlying chronic MP using the AIMP model. By combining electrophysiology, chemogenetics, and *in vivo* Ca^2+^ imaging, we demonstrated CeA-SST neurons, which are believed to be antinociceptive neurons, become hyperexcitable after the development of MP. Consistent with this finding, selective chemogenetic inactivation of CeA-SST neurons or suppression of synaptic transmission of the PBN to CeA-SST neurons alleviated chronic MP and negative emotions.

## Results

### Local application of PGB in the CeA alleviated pain in a mouse model of chronic MP

AIMP in rats or mice is considered as a preclinical model of fibromyalgia (Cheng et al., 2011; Min et al., 2011; Sluka et al., 2001). In this study, we induced chronic MP in mice using a protocol for AIMP. Mice with acidic (pH 4.0) saline injected into the gastrocnemius muscle unilaterally on day 0 (baseline, BL) and day 3 (Figure 1A) are referred to as MP mice. After the first acidic saline injection, MP mice showed a transient decrease in the paw withdrawal (PW) threshold in both ipsilateral and contralateral hind limbs in response to the von Frey filament stimulation (Figure 1B). The acute pain completely recovered by day 3 (Figure 1B). A second injection, however, caused a sustained decrease in the PW threshold that lasted for at least 14 days (Figure 1B). In contrast to the MP mice, we found that mice injected with neutral (pH 7.2) saline, referred to as the control (Ctrl) mice, showed no significant changes in the PW threshold in response to the von Frey filament stimulation (Figure 1B). Collectively, MP mice, but not Ctrl mice exhibited an increase in the PW response bilaterally that had lasted for at least two weeks (Figure 1C).

**Figure 1.**
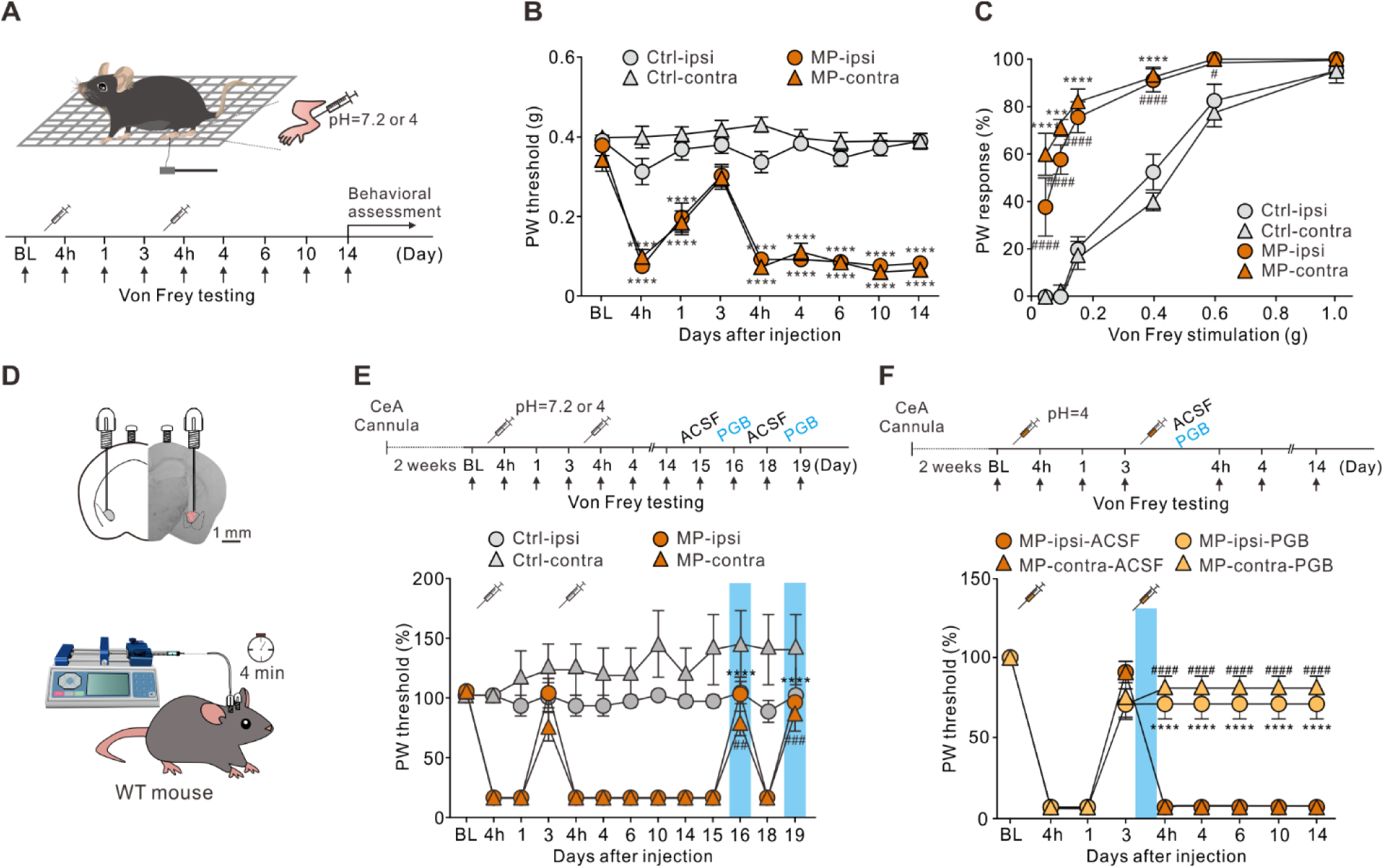
Local application of PGB in the CeA alleviated pain in a mouse model of MP. **(A)** Schematic of MP induction protocol and experimental timeline. Mice were injected with either neutral (pH 7.2, Ctrl mice) or acidic (pH 4.0, MP mice) saline into the gastrocnemius muscle unilaterally on days 0 and 3. **(B)** Plot of the PW threshold to mechanical stimuli over time (Ctrl, n = 26; MP, n = 28; two-way ANOVA with Tukey’s post hoc test, F(8,936) = 30.97, \*\*\*\**p* < 0.0001 relative to BL). **(C)** The bilateral PW responses to different filaments on day 14 in Ctrl and MP mice (Ctrl, n = 8; MP, n = 9; two-way ANOVA with Tukey’s post hoc test, F(3,180) = 121.1, ^#^*p* < 0.05, ****^,####^*p* < 0.0001; * indicates the comparison between ipsilateral hindpaws; ^#^ indicates the comparison between contralateral hindpaws). **(D)** Representative image of PGB infusion site and experimental schematic. **(E)** Effects of PGB treatment on PW threshold in Ctrl and MP mice (Ctrl, n = 9; MP, n = 9; two-way ANOVA with Tukey’s post hoc test, F(12,416) = 13.24, ^##^*p* < 0.01, ^###^*p* < 0.001, \*\*\*\**p* < 0.0001 relative to day 15; * indicates the comparison between ipsilateral hindpaws; ^#^ indicates the comparison between contralateral hindpaws. The blue area indicates the period of PGB treatment). **(F)** Top, experimental timeline. Right after the second injection of the acidic saline, one of the mouse groups was infused with PGB, and the other with ACSF. Bottom, effects of PGB treatment on PW threshold in ACSF and PGB groups (ACSF, n = 5; PGB, n = 6; two-way ANOVA with Tukey’s post hoc test, F(3,162) = 114, ****^,####^*p* < 0.001; * indicates the comparison between ipsilateral hindpaws; ^#^ indicates the comparison between contralateral hindpaws. The blue area indicates the period of PGB treatment). The following source data and figure supplement(s) for Figure 1: **Source data 1.** Numerical data to support graphs in Figure 1. **Figure supplement 1.** Comorbid affective symptoms in a mouse model of MP. **Figure supplement 1-Source data 1.** Numerical data to support graphs in Figure 1-Figure supplement 1.

In addition to mechanical allodynia, mice with chronic MP showed a variety of affective symptoms (Figure 1 - Figure supplement 1). The elevated plus maze (EPM) and light/dark (L/D) box are common approach-avoidance conflict tests based on the general aversion of mice to bright and open environments (Calhoon and Tye, 2015; Walf and Frye, 2007). Anxious mice prefer to stay in the closed arms and the dark compartment. Compared to Ctrl mice, MP mice spent less time in the open arms (Figure 1 - Figure supplement 1A) and made fewer transitions between the light and dark zones in the L/D box test (Figure 1 - Figure supplement 1B). Neither of these behavioral phenotypes was due to reduced locomotion because the total travel distances of MP and Ctrl mice were not significantly different (Figure 1 - Figure supplement 1A, B). We also tested mice in the marble burying test where anxious animals directed their energies toward minimizing threatening stimuli and tended to bury more marbles. Consistent with this notion, MP mice buried more marbles than Ctrl mice (Figure 1 - Figure supplement 1C).

Anxiety disorders are often associated with depression-like behavior and social avoidance (Allsop et al., 2014). The forced swim test (FST) was commonly used for testing depression-like behavior in mice. Compared to Ctrl mice, MP mice spent more time in the floating state, suggesting enhanced depression-like behavior (Figure 1 - Figure supplement 1D). Finally, we tested the sociability of the mice by measuring the time that mice spent in a social chamber containing a novel mouse (Yang et al., 2011). Accordingly, MP mice spent less time in the social chamber than did Ctrl mice (Figure 1 - Figure supplement 1E). The total travel distance of MP mice was not significantly different from that of Ctrl mice (Figure 1 - Figure supplement 1E). Taken together, these results suggest that MP mice suffer from mechanical allodynia and hyperalgesia and exhibit a variety of behavioral traits commonly associated with anxiety and depression. Taken together, the AIMP model satisfies the face validity of chronic MP (Calhoon and Tye, 2015).

Then, we tested the predictive validity of this model. Pregabalin (PGB) is an FDA-approved drug for treatment of chronic pain disorders and affective comorbidities. PGB is a selective ligand for α2δ subunit of voltage-gated calcium channels (VGCCs) (Häuser et al., 2009). However, how pregabalin exerts its action on both sensory and affective dimensions is unknown. The CeA receives a direct nociceptive projection from the parabrachial nucleus (PBN), which is enriched with the calcium channel α2δ subunit (Cole et al., 2005; Ikeda et al., 2007). To test whether PGB acted on the PBN-CeA pathway, we infused PGB into the CeA of Ctrl and MP mice bilaterally through a cannula (Figure 1D). Compared to Ctrl mice, cannula infusion of PGB (1 mM, 0.15 μL/site), instead of vehicle (artificial cerebrospinal fluid, ACSF), into the CeA of MP mice on day 16 and day 19 increased the PW threshold (Figure 1E). This result supports the predictive validity of this model.

In this model, the first acidic saline injection initiates a priming phase, which is reported to last for 5 to 8 days (Chen et al., 2014; Sun and Chen, 2016). Therefore, we asked whether earlier administration of PGB can prevent the development of chronic pain. To this end, we infused ACSF or PGB into the CeA of MP mice bilaterally through a cannula on day 3 right after the second acidic saline injection (Figure 1F). The group infused with ACSF was able to develop chronic pain effectively. In contrast, the pain response of the group infused with PGB became weakened (Figure 1F). This result demonstrated that local infusion of PGB into the CeA during hyperalgesic priming is sufficient to prevent the development of chronic pain.

### Local infusion of PGB into the CeA alleviated negative emotion

To test whether local PGB administration in the CeA also alleviate negative emotion in MP mice, we infused PGB into the CeA of Ctrl and MP mice bilaterally through a cannula (Figure 2A). Compared to Ctrl mice, cannula infusion of PGB (1 mM, 0.15 μL/site), instead of vehicle (artificial cerebrospinal fluid, ACSF), into the CeA of MP mice increased the open-arm time (Figure 2B) in the EPM test and decreased the number of buried marbles in the marble burying test (Figure 2C). In addition, we tested depression-like behavior and sociability of the mice using the FST and three-chambered social test. Compared to Ctrl mice, local infusion of PGB into the CeA of MP mice decreased immobility in the FST (Figure 2D) and increased the social-zone time in the three-chambered social test (Figure 2E). Notably, PGB at the dosage used in this study had no effect on locomotion (Figure 2A, E) and had little effect on affective behaviors in Ctrl mice (Figure 2B-E). These results indicated that local infusion of PGB in the CeA, in the chronic pain state, alleviated negative emotions.

**Figure 2.**
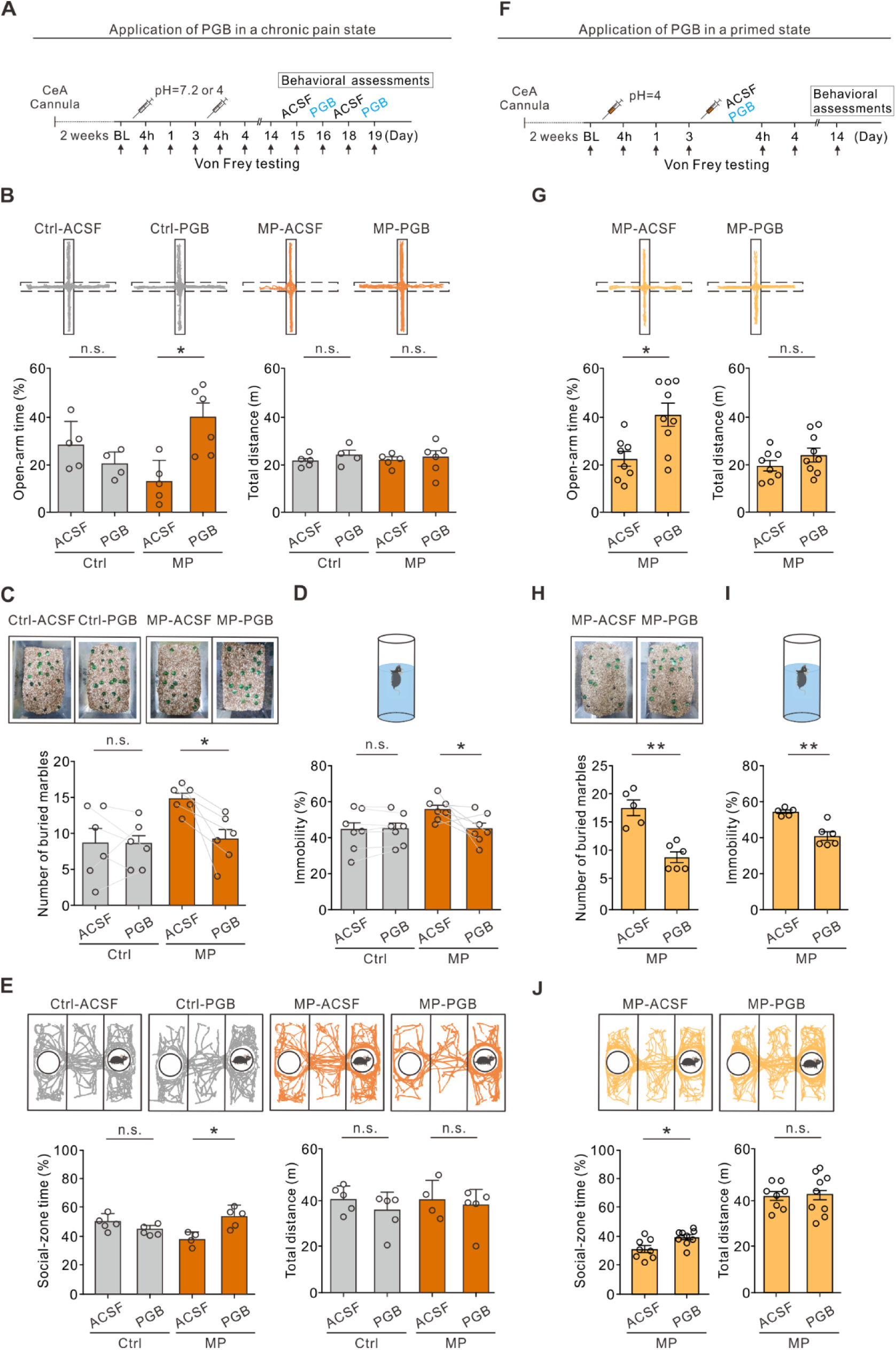
Local infusion of PGB in the CeA alleviated negative emotion. **(A)** Experimental timeline. **(B)** Top, representative travel trajectories of each Ctrl and MP group during the EPM test. Bottom, summary of the effects of PGB treatment on open-arm time (Ctrl: ACSF, 29.7 ± 6.1%, n = 5; PGB, 21.2 ± 3.6%, n = 4; Mann-Whitney test, U = 4, n.s., non-significant, *p* = 0.19. MP: ACSF, 13.8 ± 5.4%, n = 5; PGB, 39.89 ± 5.7%, n = 6; Mann-Whitney test, U = 2, \**p* = 0.017) and total distance (Ctrl: ACSF, 21.5 ± 1.7 m, n = 5; PGB, 24.1 ± 2.5 m, n = 4; Mann-Whitney test, U = 6, n.s., non-significant, *p* = 0.413. MP: ACSF, 22.0 ± 1.6 m, n = 5; PGB, 22.8 ± 2.7 m, n = 6; Mann-Whitney test, U = 11, n.s., non-significant, *p* = 0.515). **(C)** Summary of the effects of PGB treatment on the number of buried marbles (Ctrl: ACSF, 8.8 ± 2.0, n = 6; PGB, 8.5 ± 1.3, n = 6; Wilcoxon matched-pairs signed rank test, n.s., non-significant, *p* = 0.906. MP: ACSF, 14.8 ± 0.7, n = 6; PGB, 9.2 ± 1.3, n = 6; Wilcoxon matched-pairs signed rank test, \**p* = 0.031). **(D)** Summary of relative time of immobility in the FST (Ctrl: ACSF, 44.0 ± 4.3%, n = 7; PGB, 44.8 ± 3.2%, n = 7; Wilcoxon matched-pairs signed rank test, n.s., non-significant, *p* = 0.813. MP: ACSF, 56.2 ± 2.4%, n = 7; PGB, 45.4 ± 3.1%, n = 7; Wilcoxon matched-pairs signed rank test, \**p* = 0.047). **(E)** Top, representative travel trajectories of each Ctrl and MP group during the three-chamber sociability test. Bottom, summary of the effects of PGB treatment on social-zone time (Ctrl: ACSF, 49.0 ± 3.0%, n = 5; PGB, 43.4 ± 1.6%, n = 5; Mann-Whitney test, U = 4, n.s., non-significant, *p* = 0.057. MP: ACSF, 37.7 ± 2.8%, n = 4; PGB, 52.8 ± 4.2%, n = 5; Mann-Whitney test, U = 0, \**p* = 0.029) and total distance (Ctrl: ACSF, 41.3 ± 3.2 m, n = 5; PGB, 34.5 ± 4.7 m, n = 5; Mann-Whitney test, U = 6, n.s., non-significant, *p* = 0.229. MP: ACSF, 40.9 ± 4.9 m, n = 4; PGB, 36.7 ± 5.3 m, n = 5; Mann-Whitney test, U = 8, n.s., non-significant, *p* = 0.657). **(F)** Experimental timeline. Right after the second injection of the acidic saline, one of the mouse groups was infused with PGB, and the other with ACSF. The behavior assessments were performed on day 14 **(G)** Top, representative travel trajectories of each ACSF and PGB group during the EPM test. Bottom, summary of the effects of PGB treatment on open-arm time (ACSF, 22.5 ± 5.4%, n = 8; PGB, 38.6 ± 8.2%, n = 9; Mann-Whitney test, U = 10, \**p* = 0.011) and total distance (ACSF, 18.9 ± 3.2, n = 8; PGB, 26.6 ± 5.1, n = 9; Mann-Whitney test, U = 25, n.s., non-significant, *p* = 0.315). **(H)** Summary of the effects of PGB treatment on the number of buried marbles (ACSF, 17.5 ± 2.0, n = 5; PGB, 9.0 ± 1.2, n = 6; Mann-Whitney test, U = 0, \*\**p* = 0.004). **(I)** Summary of the effects of PGB treatment on the immobility time (ACSF, 54.1 ± 0.9, n = 5; PGB, 41.0 ± 4.0, n = 6; Mann-Whitney test, U = 0, \*\**p* = 0.004). **(J)** Top, representative travel trajectories of each ACSF and PGB group during the three-chamber sociability test. Bottom, summary of the effects of PGB treatment on social-zone time (ACSF, 33.4 ± 4.6, n = 8; PGB, 42.7 ± 3.3, n = 9; Mann-Whitney test, U = 11, \**p* = 0.015) and total distance (ACSF, 40.9 ± 2.6, n = 8; PGB, 41.5 ± 5.0, n = 9; Mann-Whitney test, U = 33, n.s., non-significant, *p* = 0.791). The following source data and figure supplement(s) for Figure 2: **Source data 1.** Numerical data to support graphs in Figure 2.

Given that the PGB treatment during the priming phase is sufficient to prevent the development of chronic pain, we also tested whether PGB given during the priming phase can prevent comorbid affective symptoms. To this end, we infused ACSF or PGB into the CeA of MP mice bilaterally through a cannula right after the second acidic saline injection (Figure 2F). The same behavior assessments were performed in this experiment on day 14 (Figure 2F). Compared to ACSF group, cannula infusion of PGB in the primed state increased MP mice exploration time in open arms (Figure 2G), buried less marbles (Figure 2H), decreased the immobility time (Figure 2I), and increased the social-zone time (Figure 2J). Taken together, the infusion of PGB immediately after the second acidic saline injection disrupted the hyperalgesic priming and therefore prevented the development of chronic pain-related comorbid affective emotions in MP mice.

### Chronic MP was associated with enhanced synaptic transmission and neuronal excitability in the CeA-SST neurons

To characterize maladaptive changes in amygdala circuits in chronic pain states, we recorded excitatory synaptic transmission to CeA-SST and CeA-PKCδ neurons using whole-cell patch-clamp recording in brain slices from Ctrl and MP mice (Figure 3A). Using the Cre-loxP recombination approach, we identified CeA-SST and CeA-PKCδ neurons from SST-Cre;Ai14 and PKCδ-Cre;Ai14 mice, respectively (Figure 3B). Compared to Ctrl mice, an increase in the mean frequency of spontaneous excitatory postsynaptic currents (sEPSCs) was detected in CeA-SST neurons of MP mice (Figure 3C). The mean amplitude of sEPSCs in MP mice, however, was not significantly different from that in Ctrl mice (Figure 3C). In contrast to CeA-SST neurons, recordings from CeA-PKCδ neurons of MP mice showed a significant decrease in the sEPSC frequency, but showed no change in the sEPSC amplitude compared to the Ctrl mice (Figure 3D). Similar results were observed in miniature EPSCs (mEPSCs; Figure 3 - Figure supplement 1A, B), suggesting that the transmission of excitatory inputs onto CeA-SST neurons were strengthened, while that of excitatory inputs onto CeA-PKCδ neurons were weakened.

**Figure 3.**
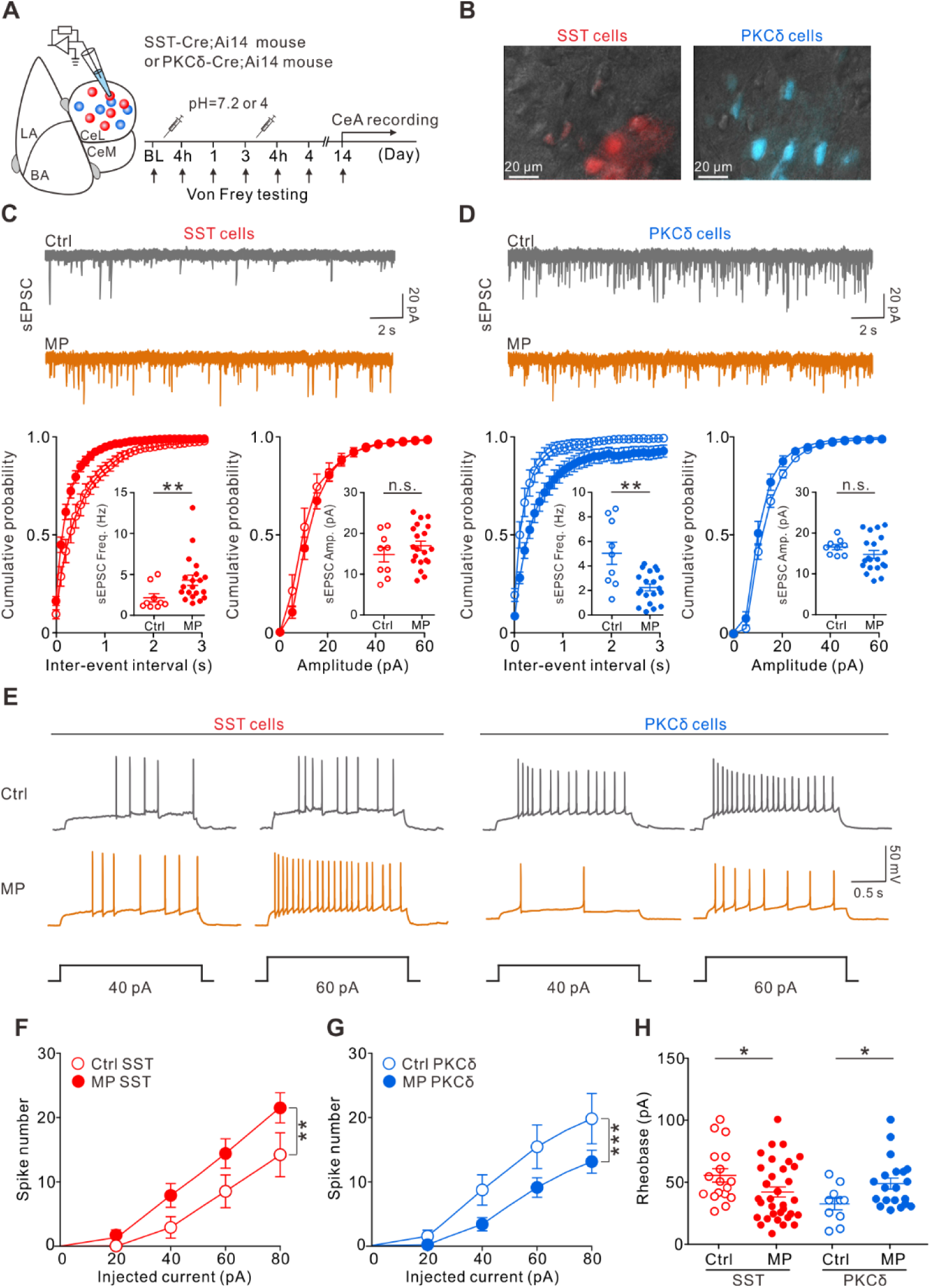
Altered excitatory transmission to CeA neurons and changed CeA neuron excitability in MP mice. **(A)**Experimental schematic and timeline. **(B)** Overlay of epifluorescence and IR-DIC images showing SST and PKCδ neurons in the CeA. Sections from SST-Cre;Ai14 and PKCδ-Cre;Ai14 mouse brains. **(C)** Top, representative sEPSC traces recorded from CeA-SST neurons of Ctrl and MP mice. Bottom, cumulative probability of inter-event interval (Ctrl, n = 9; MP, n = 20; Kolmogorov-Smirnov test, \*\**p* = 0.0014. Inset, summary of sEPSC frequency, Ctrl, 2.2 ± 0.5 Hz, n = 9; MP, 4.3 ± 0.6 Hz, n = 20; Mann-Whitney test, U = 33, \*\**p* = 0.006) and amplitude (Ctrl, n = 9; MP, n = 20; Kolmogorov-Smirnov test, n.s., non-significant, *p* = 0.998. Inset, summary of sEPSC amplitude, Ctrl, 14.9 ± 1.8 pA, n = 9; MP, 16.9 ± 1.1 pA, n = 20; Mann-Whitney test, U = 72, n.s., non-significant, *p* = 0.404). **(D)** Top, representative sEPSC traces recorded from CeA-PKCδ neurons of Ctrl and MP mice. Bottom, cumulative probability of inter-event interval (Ctrl, n = 9; MP, n = 20; Kolmogorov-Smirnov test, \*\*\*\**p* < 0.0001. Inset, summary of sEPSC frequency, Ctrl, 5.1 ± 0.9 Hz, n = 9; MP, 2.3 ± 0.3 Hz, n = 20; Mann-Whitney test, U = 36, \*\**p* = 0.0097) and amplitude (Ctrl, n = 9; MP, n = 20; Kolmogorov-Smirnov test, n.s., non-significant, *p* = 0.996. Inset, summary of sEPSC amplitude, Ctrl, 16.5 ± 0.6 pA, n = 9; MP, 14.7 ± 1.0 pA, n = 20, Mann-Whitney test, U = 60, n.s., non-significant, *p* = 0.167). **(E)** Representative responses of CeA-SST and CeA-PKCδ neurons in Ctrl and MP mice to depolarizing current injections. **(F)** Plot of number of spikes in CeA-SST neurons against injected current (Ctrl, n = 15; MP, n = 28; two-way ANOVA with Bonferroni’s multiple comparisons test, F(1,202) = 10.49, \*\**p* = 0.0014). **(G)** Plot of number of spikes in CeA-PKCδ neurons against injected current (Ctrl, n = 13; MP, n = 23; two-way ANOVA with Bonferroni’s multiple comparisons test, F(1,168) = 13.31, \*\*\**p* = 0.0004). **(H)** Summary of rheobase of CeA-SST neurons (Ctrl, 54.9 ± 5.4 pA, n = 17; MP, 41.6 ± 4.1 pA, n = 33; Mann-Whitney test, U = 178, \**p* = 0.035) and CeA-PKCδ neurons (Ctrl, 32 ± 4.7 pA, n = 10; MP, 48.5 ± 4.4 pA, n = 20; Mann-Whitney test, U = 53, \**p* = 0.038). The following source data and figure supplement(s) for Figure 3: **Source data 1.** Numerical data to support graphs in Figure 3. **Figure supplement 1.** Comparison of mEPSC, resting potential, and input resistance of CeA neurons in Ctrl and MP mice. **Figure supplement 1-Source data 1.** Numerical data to support graphs in Figure 3-Figure supplement 1.

Next, we investigated whether the intrinsic neuronal excitability of CeA-SST and CeA-PKCδ neurons was changed in MP mice (Figure 3E). To determine the changes in intrinsic properties, CeA neurons were recorded in the presence of synaptic blockers. Compared to Ctrl mice, resting membrane potentials and input resistance of CeA-SST neurons and CeA-PKCδ neurons in MP mice were not significantly altered (Figure 3 - Figure supplement 1C-F). However, CeA-SST neurons in MP mice generated more spikes in response to prolonged current injections (Figure 3E left, F). Conversely, CeA-PKCδ neurons in MP mice generated fewer spikes (Figure 3E right, G). Consistent with these results, CeA-SST neurons exhibited a lower rheobase (Figure 3H), while CeA-PKCδ neurons exhibited a higher rheobase in MP mice compared to Ctrl mice (Figure 3H). These results indicated that MP was associated with increased excitability in CeA-SST neurons and decreased excitability in CeA-PKCδ neurons.

### Inhibition of CeA-SST neurons alleviated pain and affective symptoms

Given the association between the increased CeA-SST excitability and chronic pain phenotypes in MP mice, we sought to determine the causal role of CeA-SST neurons. We therefore asked if heightened activity of CeA-SST neurons during the initial muscle injuries was required for the development of chronic pain states. To this end, a Cre-dependent AAV5 encoding an inhibitory designer receptor (hM4Di) was injected bilaterally into the CeA of SST-Cre mice (Figure 4A). Whole-cell current-clamp recordings from acute amygdala slices prepared from mice expressing hM4Di in the CeA showed that bath application of clozapine-n-oxide (CNO; 5 µM) selectively inhibited spontaneous firing in hM4Di-expressing SST neurons (Figure 4B, C). To test the essential role of CeA-SST neuronal activity in MP induction, CNO (5 mg/kg body weight) was intraperitoneally (i.p.) injected into mice immediately after the second acidic saline injection. Compared to mCherry-MP mice, silencing CeA-SST neurons during primed state in hM4Di-MP mice prevented the development of MP (Figure 4D). This result suggests that activation of CeA-SST neurons after peripheral injury was necessary for pain chronification.

**Figure 4.**
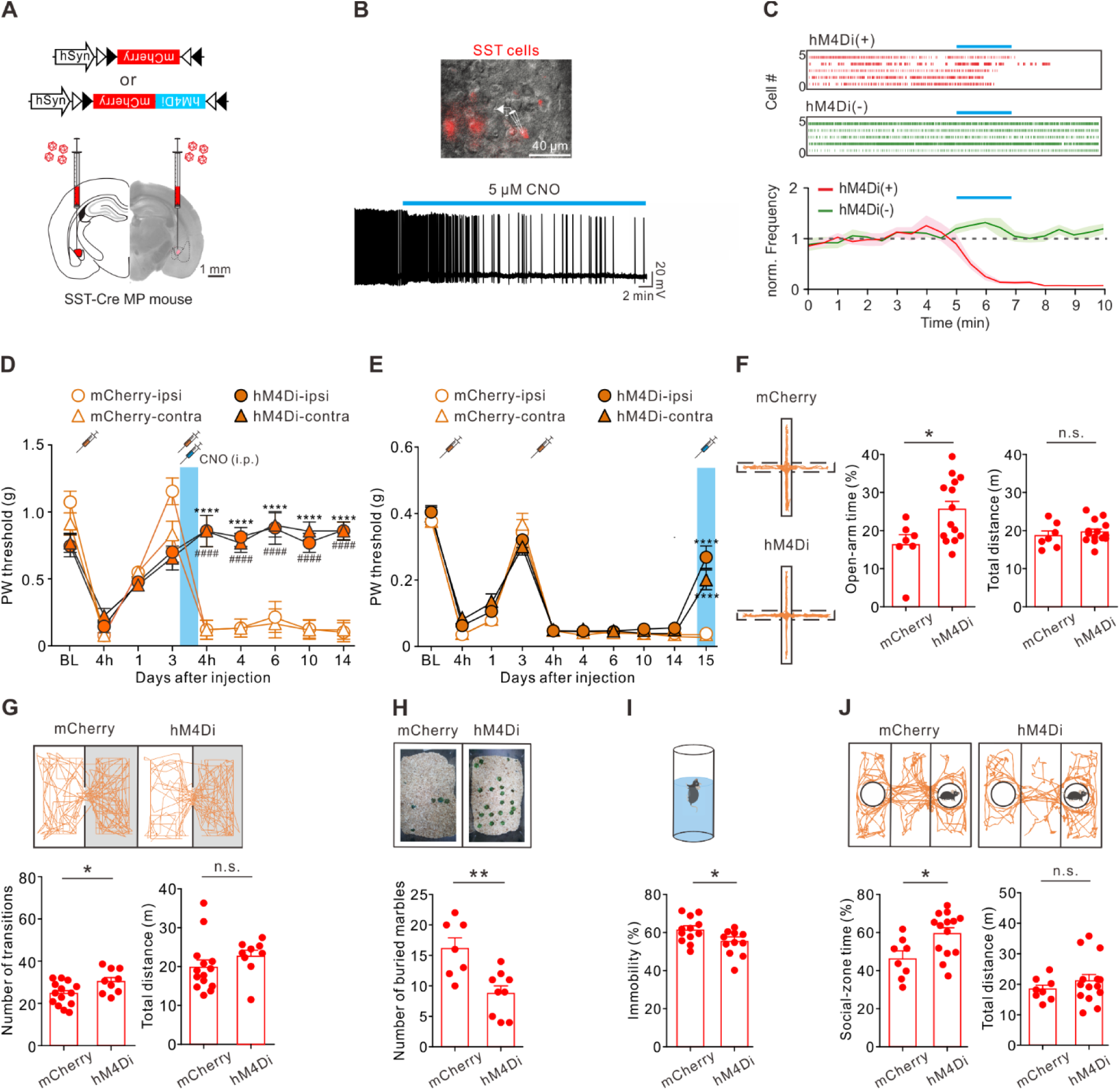
Inactivating SST neurons alleviated pain and affective symptoms. **(A)** Viral constructs and experimental schematic. **(B)** Top, overlay of epifluorescence and IR-DIC images showing mCherry(+) neurons in the CeA. Bottom, membrane potential changes of an hM4Di-expressing CeA-SST neuron before and after bath application of CNO. **(C)** Top, representative of raster plots of hM4Di(+) and hM4Di(-) neurons. Blue bar indicates 5 µM CNO application for 2 min. Bottom, normalized firing frequency in both hM4Di(+) and hM4Di(-) neurons (n = 10 neurons for each group). **(D)** Effects of CNO treatment on day 3 on PW threshold (mCherry, n = 5; hM4Di, n = 9; two-way ANOVA with Tukey’s post hoc test, F(3,216) = 46.82, ****,^####^*p* < 0.0001; * indicates the comparison between ipsilateral hindpaw; ^#^ indicates the comparison between contralateral hindpaws. The blue area indicates the period of CNO treatment). **(E)** Effects of CNO treatment on PW threshold in MP mice (mCherry, n = 10; hM4Di, n = 52; two-way ANOVA with Tukey’s post hoc test, F(3,1200) = 3.18, \*\*\*\**p* < 0.0001 relative to day 14. The blue area indicates the period of CNO treatment). **(F)** Left, representative trajectories of each MP group during the EPM test. Right, summary of the effects of CNO treatment on open-arm time (mCherry, 16.2 ± 2.5%, n = 7; hM4Di, 25.5 ± 2.2%, n = 14; Mann-Whitney test, U = 19, \**p* = 0.025) and total distance (mCherry, 18.7 ± 1.3 m, n = 7; hM4Di, 19.7 ± 0.8 m, n = 14; Mann-Whitney test, U = 37, n.s., non-significant, *p* = 0.383). **(G)** Top, representative travel paths of each MP group during the L/D box test. Bottom, summary of CNO effect on the number of transitions (mCherry, 23.6 ± 1.5, n = 14; hM4Di, 30.0 ± 1.9, n = 9; Mann-Whitney test, U = 28.5, \**p* = 0.028) and total distance (mCherry, 20.0 ± 1.8 m, n = 14; hM4Di, 22.7 ± 1.6 m, n = 9; Mann-Whitney test, U = 34, n.s., non-significant, *p* = 0.072). **(H)** Top, representative images of the marble burying test. Bottom, summary of the effects of CNO treatment on the number of buried marbles (mCherry, 16.1 ± 1.8, n = 7; hM4Di, 8.8 ± 1.2, n = 9; Mann-Whitney test, U = 6.5, \*\**p* = 0.005). **(I)** Top, schematic of the FST setup. Bottom, summary of relative time of immobility (mCherry, 61.4 ± 1.9%, n = 12; hM4Di, 54.2 ± 2.0%, n = 11; Mann-Whitney test, U = 29, \**p* = 0.022). **(J)** Top, representative travel paths of each MP group during the three-chamber sociability test. Bottom, summary of the effects of CNO treatment on social-zone time (mCherry, 46.7 ± 3.7%, n = 8; hM4Di, 60.0 ± 3.1%, n = 14; Mann-Whitney test, U = 22, \**p* = 0.019) and total distance (mCherry, 18.5 ± 1.3 m, n = 8; hM4Di, 21.2 ± 2.1 m, n = 14; Mann-Whitney test, U = 47, n.s., non-significant, *p* = 0.55). The following source data and figure supplement(s) for Figure 4: **Source data 1.** Numerical data to support graphs in Figure 4. **Figure supplement 1.** Suppressing CeA-SST neuron excitability in Ctrl mice exerted little effect on nociception and affective behaviors. **Figure supplement 2.** Enhancing PKCδ neuron excitability reduced pain, but failed to alleviate affective symptoms. **Figure supplement 3.** Proposed wiring diagram of CeA circuits, extrinsic pathways and behavioral responses. **Figure supplement 1-Source data 1.** Numerical data to support graphs in Figure 4-Figure supplement 1. **Figure supplement 2-Source data 1.** Numerical data to support graphs in Figure 4-Figure supplement 2.

Next, we test the causal role of CeA-SST neurons in MP-associated behaviors. CNO was i.p. injected into mice after MP had been established (Figure 4E). We found that inhibiting CeA-SST neurons in MP mice 2 weeks after MP induction was still effective in increasing the PW threshold on day 15 (Figure 4E), and the pain-reduction effect lasted for 24 hours and persisted if CNO was given repeatedly (data not shown). On the same day following CNO injection, we also tested the effect of inhibiting CeA-SST neurons on the affective symptoms. Compared to mCherry-MP mice, hM4Di-MP mice spent more time in open arms during the EPM test (Figure 4F), exhibited increased transitions between light and dark zones in the L/D box test (Figure 4G), buried fewer marbles in the marble burying test (Figure 4H), and immobilized less in the FST (Figure 4I). Moreover, hM4Di-MP mice increased the time spent in the social zone in the three-chamber sociability test (Figure 4J). However, the total travel distances of mCherry-MP and hM4Di-MP mice were not significantly different (Figure 4F, G, J). Notably, chemogenetic inhibition of CeA-SST neurons in hM4Di-Ctrl mice had little effects on the PW threshold and on their respective performances in the EPM, L/D box, marble burying test, FST, and sociability test (Figure 4 - Figure supplement 1). Together, these observations suggest that inactivating CeA-SST neurons selectively reduced pain and alleviated anxiety- and depression-like behaviors in MP mice.

Both enhanced CeA-SST neuronal activity and decreased CeA-PKCδ neuronal activity were observed in MP mice. Moreover, these two types of neurons reciprocally inhibit each other. Next, we tested whether increasing CeA-PKCδ neuronal activity could also reduce mechanical hyperalgesia and anxiety-like behavior. To this end, the excitatory designer receptor (hM3Dq) was virally expressed in CeA-PKCδ neurons (Figure 4 - Figure supplement 2A). Similar to the silencing of hM4Di-expressing CeA-SST neurons, the activation of hM3Dq-expressing CeA-PKCδ neurons greatly increased the PW threshold in MP mice (Figure 4 - Figure supplement 2B). Intriguingly, unlike the results of inactivating CeA-SST neurons, enhancing CeA-PKCδ neurons had little effects on anxiety-like behaviors. There were no changes in the open-arm time (Figure 4 - Figure supplement 2C), number of transitions during the L/D box test (Figure 4 - Figure supplement 2D), and number of buried marbles (Figure 4 - Figure supplement 2E). Similarly, enhancing CeA-PKCδ neurons did not improve depression-like behavior (Figure 4 - Figure supplement 2F) and sociability (Figure 4 - Figure supplement 2G). Taken together, these results suggest that activation of CeA-PKCδ neurons is sufficient to reduce mechanical hyperalgesia in MP mice. However, changes in the excitability of CeA-PKCδ neurons are not causally related to chronic MP-related affective behaviors.

### PGB suppressed CeA-SST neuron excitability and nociceptive transmission onto CeA-SST neurons

Since enhanced CeA-SST neuronal activity were observed in MP mice, we next tested the effect of PGB on excitatory synaptic transmission onto CeA-SST neurons. To this end, we injected an AAV5-CaMKIIα-ChR2-eYFP virus into the PBN of SST-Cre;Ai14 mice which were then established into either a Ctrl or a MP mouse four weeks later (Figure 5A). Using the optogenetic stimulation, we recorded light-evoked EPSCs from CeA-SST neurons of Ctrl or MP mice in response to short light (470 nm, 5 ms) illumination to PBN terminals in the CeA region (Figure 5B; Ctrl, left; MP, right). Notably, bath application of PGB decreased the amplitude of light-evoked EPSCs in both Ctrl and MP mice (Figure 5C, D). The reduction of EPSCs was concomitant with an increase in the paired-pulse ratio (interpulse interval = 200 ms; Figure 5E).

**Figure 5.**
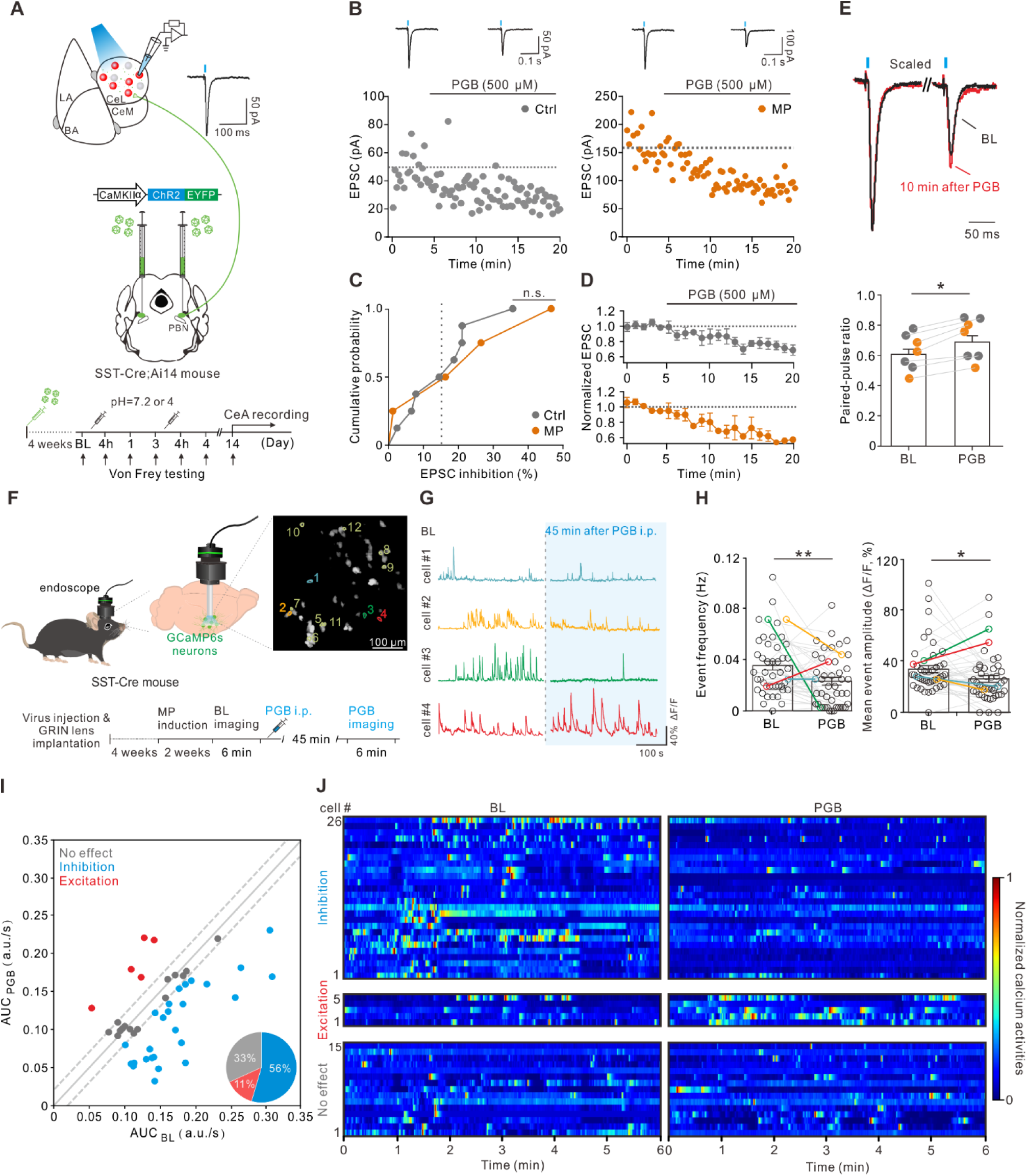
PGB suppressed CeA-SST neuron excitability and glutamate release from PBN to CeA-SST neurons. **(A)** Experimental schematic and timeline. **(B)** Left, amplitude of the EPSC before and after PGB application of CeA-SST neurons in Ctrl mice. Right, amplitude of the EPSC before and after PGB application of CeA-SST neurons in MP mice; traces of EPSCs of the cell are shown above. **(C)** The cumulative probability of EPSC inhibition of Ctrl and MP group (Ctrl, n = 8; MP, n = 4; dashed line: 15% EPSC inhibition). **(D)** Cells with EPSC inhibition greater than 15% are pooled and plotted for the normalized EPSC over time (Ctrl, n = 4; MP, n = 3). **(E)** Top, representative EPSC traces before and 10 min after PGB application. Average trace after PGB application is normalized to the peak of average EPSC trace during BL. Paired-pulse interval = 200 ms. Bottom, summary of paired-pulse ratio before and after PGB application (BL, 0.60 ± 0.04, n = 7; PGB, 0.68 ± 0.05, n = 7; gray circle: Ctrl group; orange circle: MP group; Wilcoxon matched-pairs signed rank test, \**p* = 0.016). **(F)** Left and bottom, experimental schematic and timeline. Right, representative image of CeA-SST neurons labeled with GCaMP6s. Cells with their calcium traces shown in the panel G were labeled. **(G)** Representative traces from cells of interest (color-matched in the panel a right). **(H)** The frequency of calcium events for CeA-SST neurons before and after PGB i.p. injection (BL, 0.04 ± 0.003 Hz; PGB, 0.02 ± 0.003 Hz, n = 46 cells from 3 mice; Wilcoxon matched-pairs signed rank test, \*\**p* = 0.004) and the percentage of ΔF/F before and after PGB i.p. injection (BL, 34.1 ± 2.8 %; PGB, 26.4 ± 2.7 %, n = 46 cells from 3 mice; Wilcoxon matched-pairs signed rank test, \**p* = 0.045). **(I)** The AUC of Ca^2+^ traces of CeA-SST neurons before and after PGB application. Neurons with ΔAUC < 1 × *σ_dev_* (0.02 a.u./s) were distributed within two dashed lines. Pie chart, the percentage of neuronal responses after PGB treatment (n = 46 cells). **(J)** The heatmaps of normalized Ca^2+^ activities before and after the PGB treatment. Neurons were grouped based on the classification in (I). The following source data and figure supplement(s) for Figure 5: **Source data 1.** Numerical data to support graphs in Figure 5.

Finally, we tested whether i.p. injection of PGB suppresses CeA-SST neuron activity using the *in vivo* calcium imaging. To selectively image the calcium activities of CeA-SST neurons, SST-Cre mice were virally expressed with the calcium indicator GCaMP6s and implanted with the GRIN lens above the CeA for four weeks before the test. Calcium activities of single CeA-SST neurons were detected before and 45 min after PGB i.p. injection (30 mg/kg; Figure 5F), which is known to significantly increase the mechanical threshold of hindpaws in this MP model (Yokoyama et al., 2007). After PGB i.p. injection, the CeA-SST neurons exhibited decreased calcium event frequency (Figure 5G, H left) and mean amplitude (ΔF/F) (Figure 5G, H right). The area under curve (AUC) of spontaneous Ca^2+^ activities suggested that in the presence of PGB, approximately 56% (26/46 cells) CeA-SST neurons were inhibited, 11% (5/46 cells) CeA-SST neurons were excited, and 33% (15/46 cells) CeA-SST neurons were insensitive to PGB application (Figure 5I, J). Taken together, these results suggest that PGB suppressed synaptic transmission of glutamate onto CeA-SST neurons, thereby reducing mechanical allodynia and reversed anxiety- and depression-like behaviors in MP mice.

## Discussion

In this study, we established a chronic MP model in mice. After the first acidic saline injection, the CeA neurons were transformed into the primed state. The second acidic saline injection during the hyperalgesic priming promoted pain chronification. Local application of PGB into the CeA reduced mechanical allodynia and reversed affective behaviors in chronic MP mice. Intriguingly, intra-CeA application of PGB immediately after the second acidic saline prevented pain chronification. Notably, the chronic MP was accompanied by enhanced glutamatergic transmission onto CeA-SST neurons, and decreased synaptic transmission onto CeA-PKCδ neurons. Furthermore, CeA-SST neuron excitability was increased, whereas CeA-PKCδ neuron excitability was reduced. In agreement with the role of CeA-SST neurons in central sensitization, chemogenetic inactivation or pharmacological suppression of CeA-SST neurons by PGB effectively alleviates chronic MP and comorbid affective behaviors.

### Supraspinal mechanisms of PGB

VGCCs comprise a series of polypeptides of the principal α1 subunit, auxiliary β and α2δ subunits (Catterall, 2000; Uchitel et al., 2010). While PGB and GBP belongs to a family of GABA analog drugs called gabapentinoid, it binds to the α2δ subunit-containing calcium channels, instead of GABA receptors (Uchitel et al., 2010). In the CNS, α2δ subunits, including subtype 1 (α2δ-1) and 2 (α2δ-2) subunits, are expressed at axon terminals and are therefore potential targets of PGB (Bian et al., 2006; Cain et al., 2017; Ma et al., 2018; Uchitel et al., 2010). Previous studies show that α2δ subunit is upregulated in injured dorsal root ganglion neurons before the development of the allodynia (Luo et al., 2001; Matsuzawa et al., 2014). In addition, the mouse model of chronic migraine exhibits the gain-of-function of VGCCs (Cain et al., 2017). Furthermore, the expression of α2δ subunit diminishes in nerve-ligated animals after recovering from tactile allodynia (Luo et al., 2001). Several studies have shown that the application of PGB reduces the calcium influx at nerve terminals and in turn decreases the release of neurotransmitters by binding to α2δ subunits (Cunningham et al., 2004; Dooley et al., 2000; Matsuzawa et al., 2014). These studies supported our observation that application of PGB not only alleviated chronic MP but also prevented the development of chronic pain. It is worth noting that our study demonstrates the exact synapses and specific neuron types by which PGB exerts its action on synaptic transmission in a chronic MP model.

### Maladaptive rewiring of glutamatergic synapses onto CeA neurons

Acid-induced MP is associated with enhanced pERK expression in the PVT and CeA (Chen et al., 2010), indicating the involvement of central sensitization. Consistent with the previous study, we observed that maladaptive rewiring of CeA circuits was associated with the generation of chronic pain. Several excitatory inputs impinge onto CeA neurons (Fadok et al., 2018; Ikeda et al., 2007). The CeA receives nociceptive inputs from the dorsal horn via the PBN and affect-related information from the LA (Ikeda et al., 2007; Li and Sheets, 2020; Wilson et al., 2019). The PBN-CeA synaptic potentiation is consolidated in chronic neuropathic pain (Ikeda et al., 2007). In addition, BLA-CeA synapses are potentiated in chronic neuropathic pain (Ikeda et al., 2007). The PVT, a structure that is readily activated by both physical and psychological stressors, also projects to the CeA. The PVT has been demonstrated to be required for LTP of excitatory synaptic transmission to CeA-SST neurons (Penzo et al., 2015). Taken together, our results revealed that the net effect of excitatory synapses onto CeA-SST neurons is strengthened, whereas the respective synapses onto CeA-PKCδ neurons are weakened in chronic MP.

### Subtype of CeA neurons are sensitized in different mouse model of chronic pain

A recent study showed that neuropathic pain-induced activated forms of ERK (pERK) and cFos are preferentially expressed in CeA-PKCδ neurons, which exhibit enhanced excitability relative to CeA-SST neurons. They showed that chemogenetic silencing of CeA-PKCδ neurons or activation of CeA-SST neurons reversed nerve-injury-induced hyperalgesia (Wilson et al., 2019). Another study showed that chemogenetic inhibition of the CeA-PKCδ neurons reduces the mechanical hyperalgesia in mice with the formalin-induced pain (Chen et al., 2022). However, two recent studies contradict this notion (Hua et al., 2020; Zhou et al., 2019). Using the spared nerve injury model, Zhou and colleagues demonstrated that a small population of CeA-SST neurons, which are glutamatergic and project to the lateral habenula (LHb), exhibit enhanced excitability and are involved in a neural circuit for hyperalgesia and comorbid depressive symptoms (Zhou et al., 2019). Moreover, a subpopulation of neurons co-expressing PKCδ and enkephalin confined to the mid-posterior axis of the CeA was shown to mediate antinociceptive function in response to general anesthetics (Hua et al., 2020). In our study, using the AIMP model, we observed that chemogenetic inactivation of CeA-SST neurons effectively alleviates chronic MP and comorbid affective behaviors. Overall, the subtype of sensitized neurons in the CeA may depend on the chronic pain models.

### The CeA contains functionally distinct neuronal populations

CeA-SST neurons form reciprocal connections with CeA-PKCδ neurons (Ciocchi et al., 2010; Haubensak et al., 2010). Notably, silencing CeA-SST neurons in the chronic MP model improved affective behaviors, while activating CeA-PKCδ neurons had little effect on it. Our results support a notion that CeA-SST neurons contain heterogeneous subpopulations (Penzo et al., 2014). CeA-SST neurons may comprise at least two distinct subtypes of pro-nociceptive neurons in the chronic MP model. A subset of CeA-SST neurons may project to the BNST for approach-avoidance behavior, the midbrain vlPAG for active avoidance behavior, and the LHb for depression-like behavior. We speculate that CeA-PKCδ neurons may inhibit the other subset of CeA-SST neurons, which form reciprocal inhibition with CeA-PKCδ neurons. In agreement with this hypothesis, chemogenetic activation of CeA-PKCδ neurons reduces mechanical allodynia without influencing affective behaviors in MP. Of note, investigations on pain-related affective comorbidities at the cellular level between these two neurons have not been reported in the acid-induced MP model. Nevertheless, the neural mechanisms by which CeA-SST neurons and their downstream targets regulate negative affect have been characterized (Ahrens et al., 2018; Haubensak et al., 2010; Li et al., 2013; Zhou et al., 2019). Therefore, the hyperexcitability of CeA-SST neurons and the resulting affective symptoms can be well explained by the downstream pathways of CeA-SST neurons (Figure 4 - Figure supplement 3).

A potential caveat of this study is that single molecular markers such as SST or PKCδ may not be specific enough to identify functionally distinct neuronal populations in the modulation of pain. To achieve greater specificity, composite molecular markers and/or anatomical locations should be taken into consideration. A single molecularly defined population can be further divided based on its location within a brain area (Li and Sheets, 2020; Wilson et al., 2019). For example, spared nerve injury distinctly alters inputs from the PBN to SST neurons based on their location in the lateral division of CeA (CeL). Notably, input from the PBN to SST neurons in the capsular division of CeA (CeC) is depressed, whereas the same input to SST neurons in the CeL and the medial division of CeA (CeM) is not altered. Moreover, our recent study (Hou et al., 2016) reported that CeA-SST neurons exhibit a high degree of variation in the spike delay in response to the current injection in the slice recording and added an additional layer of heterogeneity. Given the distinct synaptic and cellular properties of these subpopulations, they are likely to react differently to chemogenetic manipulations. Thus, while the entire molecularly defined neuronal population is targeted for manipulations, the net effects are likely to be dominated by the subpopulations that are preferentially labelled. In sum, a novel toolkit by integrating anatomical, physiological, and molecular profiles of single neurons is needed for functional dissection of CeA microcircuitry.

## Materials and Methods

### Animals

Four transgenic mouse lines were used in this study: SST-Cre (stock no. 013044), SST-Cre;Ai14 (SST-Cre line crossed with Ai14 line), PKCδ-Cre (stock no. 011559), and PKCδ-Cre;Ai14 (PKCδ-Cre line crossed with Ai14 line). The SST-Cre line and Ai14 tdTomato reporter (stock no. 007914) were purchased from the Jackson Laboratory. PKCδ-Cre and C57BL/6J mice were purchased from Mutant Mouse Resource and Research Centers (MMRRC) and National Laboratory Animal Center (NLAC), respectively. Mice aged 2–5 months of either sex were used in the electrophysiological and behavioral studies. All mice were bred onto the C57BL/6J genetic background. Mice were housed in a 12-h light-dark cycle and given food and water *ad libitum*. Two-month-old mice were injected with a virus and implanted with optical fibers in the CeA. The animals were handled in accordance with national and institutional guidelines. All behavioral procedures were conducted in accordance with the protocol approved by the Institutional Animal Care and Use Committee (IACUC) of the National Yang Ming Chiao Tung University.

### Viruses

To specifically express designer receptors exclusively activated by designer drugs (DREADDs) onto CeA-SST and CeA-PKCδ neurons, we used a recombinant adeno-associated virus serotype 5 (AAV5) carrying hM3Dq or hM4Di conjugated to mCherry in a double-floxed inverted open reading frame (DIO), driven by the human Synapsin I (hSyn) promoter (AAV5-hSyn-DIO-hM3Dq-mCherry or AAV5-hSyn-DIO-hM4Di-mCherry). In the optogenetic experiments, AAV5 carrying channelrhodopsin-2 (ChR2) conjugated to eYFP driven by the CaMKIIα promotor (AAV5-CaMKIIα-ChR2-eYFP), was used to selectively express ChR2 in PBN neurons. To silence neurons, AAV5-hSyn-DIO-hM4Di-mCherry was injected into the CeA region. To activate neurons, the AAV5-hSyn-DIO-hM3Dq-mCherry virus was injected into the CeA region and the AAV5-CaMKIIα-ChR2-eYFP virus was injected into the PBN region. In addition, a viral vector carrying the red fluorescent protein (AAV5-hSyn-DIO-mCherry) was used as control. In the *in vivo* calcium experiment, the AAV5-Syn-Flex-GCaMP6s-WPRE-SV40 was injected into the CeA region. All viral vectors were purchased from the Vector Core at the University of North Carolina (Chapel Hill, NC, USA) or Addgene Vector Core (Watertown, MA, USA).

### Acid-induced muscle pain model

The model of acid-induced muscle pain (Sluka et al., 2001) was used as a preclinical fibromyalgia-like MP model. All mice were briefly anesthetized with isoflurane (4% induction, 1.5%–2% maintenance in O_2_; Halocarbon Laboratories, North Augusta, SC, USA). After anesthesia, MP and Ctrl mice received 20-μL injections of acidic saline (pH 4.0) and neutral saline (pH 7.2), respectively, on day 0 in the left gastrocnemius muscle. After three days (day 3), the same gastrocnemius muscle was re-injected with acidic or neutral saline. The pH value of the 2-(N-morpholino) ethanesulfonic acid (MES)-buffered saline (154 mM NaCl, 10 mM MES) was used to construct both the acidic and neutral saline, while the pH values were adjusted with 0.1 M HCl or NaOH.

### Immunohistochemistry

Mice were deeply anesthetized and sequentially perfused through the left ventricle with PBS (0.9% NaCl in 0.01 M phosphate buffer, pH 7.4), followed by 30 mL of ice-cold 4% paraformaldehyde in 0.1 M PBS. The brain was rapidly removed and fixed in 4% paraformaldehyde in 0.1 M PBS for 6 h at 4°C, after which it was cryoprotected with 15% sucrose in 0.1 M PBS for 24 h at 4°C, followed by 30% sucrose in 0.1 M PBS for 24 h at 4°C. Coronal brain sections (45-μm thickness) containing the amygdala and surrounding regions were cut using the cryostat microtome (CM1900, Leica Microsystems, Nussloch, Germany). Sections were treated with 3% H_2_O_2_ for 10 min and then blocked with 0.1% Triton X-100 in TBS containing 2% BSA and 2% normal goat serum (NGS, Vector Laboratories, Burlingame, CA, USA) for 2 h at room temperature. To confirm the expression of virus in the CeA, slices were stained with a rabbit anti-red fluorescent protein (RFP) primary antibody (1:300; Rockland, Limerick, PA, USA) and Alexa-594 conjugated goat anti-rabbit secondary antibody (1:500; Invitrogen, Carlsbad, CA, USA) or Alexa-488 conjugated goat anti-rabbit secondary antibody (1:500; Invitrogen, Carlsbad, CA, USA). The staining results were examined and photographed under a fluorescence microscope (BX63, Olympus, Tokyo, Japan) or a confocal laser excitation microscope (Leica SP5, Leica Microsystems, Wetzlar, Germany).

### Stereotaxic surgery

Mice were deeply anesthetized with isoflurane (4% induction, 1.5%–2% maintenance in O_2_; Halocarbon Laboratories, North Augusta, SC, USA) and placed in a stereotaxic injection frame (IVM-3000; Scientifica, Uckfield, UK). The injections were performed using the following stereotaxic coordinates: For CeA, 1.31 mm posterior from bregma, 2.87 mm lateral from the midline on both sides, and 4.72 mm ventral from the cortical surface; For PBN, 5.1 mm posterior from bregma, 1.2 mm lateral from the midline on both sides, and 3.2 mm ventral from the cortical surface. During all surgical procedures, mice were kept on a heating pad (TCAT-2LV CONTROLLER, Physitemp Instruments, Clifton, NJ, USA or Physiological Biological Temperature Controller TMP-5b, Supertech Instruments, Budapest, Hungary) to maintain their surface body temperatures at 34°C. After securing the head with ear bars, 75% ethanol was used to sterilize the surgical area, and the eyes were protected using an ophthalmic gel. For viral injections, we bilaterally injected 0.35 μL and 0.5 μL of the viral solution into the CeA and PBN, respectively, using a 10-μL NanoFil syringe and a 34-G beveled metal needle (World Precision Instruments, Sarasota, FL, USA). The flow rate (0.1 μL/min) was controlled with a nanopump controller (KD Scientific, Holliston, MA, USA). After viral injection, the needle was raised 0.1 mm above the injection site for an additional 10 min to allow the virus to diffuse before being withdrawn slowly. To reach optimal viral expression, all animals were allowed to recover for at least 4 weeks before behavioral and electrophysiological experiments.

### Cannula implantation for intra-CeA drug application

The cannula for implantation consisted of the guide cannula (27-G, 0.41 mm in diameter and 5 mm long; RWD Life Science, Shenzhen, China) and dummy cannula (0.2 mm in diameter and 5.5 mm long; RWD Life Science, Shenzhen, China). The guide cannulas were placed in the CeA (±2.87 mm lateral, 1.31 mm posterior, 4.65 mm ventral) bilaterally. To fix the guide cannulas onto the skull, dental resin cement C&B Super-Bond (Sun Medical, Moriyama, Japan) was applied to the surface of the skull around the cannula for approximately 10 min. After the resin cement hardened, the cannula was removed from the homemade holder and the mice were placed back into their home cages for recovery. One week after recovery, mice underwent behavioral tests. During the test, the dummy cannulas were replaced by internal cannulas (0.21 mm in diameter and 5.5 mm long; RWD Life Science, Shenzhen, China). We bilaterally injected 0.15 μL of artificial cerebrospinal fluid (ACSF) at a rate of 75 nL/min through the cannulas on day 15 and 18. For the delivery of PGB (Pfizer, New York, NY, USA), we bilaterally injected 0.15 μL of 1 mM PGB at a rate of 75 nL/min through the cannulas on day 16 and 19. For priming experiment, we bilaterally injected 0.15 μL of ACSF or PGB through the cannulas right after the second acidic saline injection on day 3.

### *In vivo* calcium imaging

After virus injection, the GRIN lens (diameter, 0.5 mm; length, 6.1 mm; Inscopix, CA, USA) was placed in the CeA (2.87 mm lateral, 1.31 mm posterior, 4.72 mm ventral) region. To fix the lens onto the skull, dental resin cement C&B Super-Bond (Sun Medical, Moriyama, Japan) was applied to the surface of the skull around the lens for approximately 10 min. After the resin cement hardened, the lens was removed from the holder and the mice were placed back into their home cages for recovery. Three to four weeks after virus injection, mice underwent the induction of MP. Two weeks after MP induction, the calcium activities of CeA-SST neurons were recorded. Calcium signals were detected using miniscope system (nVista 3.0; Inscopix, Palo Alto, CA, USA). The blue LED power ranged from approximately 1.4 mW at the focal plane. The images (1280 x 800 pixels) were acquired at 15 or 20 Hz frame rate. After adapting the miniscope, the calcium imaging was recorded from freely-moving mice in the home cage. The analysis was performed using the Inscopix data processing software (version 1.6.0, Inscopix, Palo Alto, CA, USA). The constrained nonnegative matrix factorization (CNMFE) was applied to extract calcium event traces (ΔF/F) of all the recorded CeA-SST neurons.

### Calcium signal analysis

Before and after PGB treatment, the frequencies and mean amplitudes of events were detected using the built-in template searching function (Clampfit 10.3; Molecular Devices, Sunnyvale, CA, USA). For the area under curve (AUC) analysis, in order to remove calcium-independent residual noise and to improve the signal-to-noise ratio, the Gaussian kernel filter 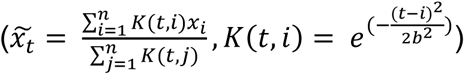 with *b* (bandwidth) set as 10 frames was applied to the whole calcium trace of single neurons. Subsequently, calcium signals (ΔF/F) before and after PGB treatment were concatenated and normalized to the range of 0 to 1 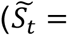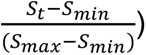. For both BL and PGB trials, calcium traces recorded in the initial 6 minutes were extracted and resampled to 10 Hz by interpolation. Spontaneous activity of a neuron was measured via calibrating its AUC per second from the normalized calcium activity of a given trial. The standard deviation of all neurons’ changes in AUC per second after PGB application (*σ_dev_*) was arbitrarily used as a cutoff threshold to identify PGB-responsive neurons. After application of PGB, neurons with ΔAUC greater than 1 × *σ_dev_* were defined as either the PGB excitation or inhibition group. Otherwise, neurons were categorized as the no effect group.

### Slice preparation and electrophysiology

After behavioral tests, AAV-injected mice were sacrificed and acute coronal brain slices of 300-μm thickness were cut using a vibratome (DTK-1000; Dosaka, Kyoto, Japan) in ice-cold sucrose saline containing the following (in mM): 87 NaCl, 25 NaHCO_3_, 1.25 NaH_2_PO_4_, 2.5 KCl, 10 glucose, 75 sucrose, 0.5 CaCl_2_, and 7 MgCl_2_. Slices were allowed to recover in an oxygenated (95% O_2_ and 5% CO_2_) sucrose saline-containing chamber at 34°C for 30 min before being maintained at room temperature until recording. During the experiment, slices were transferred to a submerged chamber and perfused with oxygenated ACSF containing the following (in mM): 125 NaCl, 25 NaHCO_3_, 1.25 NaH_2_PO_4_, 2.5 KCl, 25 glucose, 2 CaCl_2_, and 1 MgCl_2_. The expression of the virus or tdTomato expression was confirmed by red or green fluorescence and neurons in the CeA were visually selected for recordings under an infrared differential interference contrast microscope (BX51WI, Olympus, Tokyo, Japan) equipped with an LED source (505 nm and 590 nm, LED4D162, controlled by DC4104 driver, Thorlabs, Newton, NJ, USA). For optical stimulation, ChR2-expressing neurons were excited by 470 nm LED light (driven by DC4104 driver, Thorlabs, Newton, NJ, USA). Whole-cell patch-clamp recordings were conducted with an Axopatch 200B amplifier (Molecular Devices, Sunnyvale, CA, USA). Recording electrodes (3–6 MΩ) were pulled from borosilicate glass capillaries (outer diameter, 1.5 mm; 0.32 mm wall thickness; Harvard Apparatus). The glass microelectrodes were filled with a low Cl^-^ internal solution containing the following (in mM): 136.8 K-gluconate, 7.2 KCl, 0.2 EGTA, 4 MgATP, 10 HEPES, 7 NA2-phosphocreatine, 0.5 Na_3_GTP and 0.4% biocytin (wt/vol; 310 mOsm/L). The pipette capacitance was compensated. Signals were low-pass filtered at 5 kHz and sampled at 10 kHz using a digitizer (Digidata 1440A; Molecular Devices, Sunnyvale, CA, USA). In *ex vivo* slice recordings, the following antagonists were added to the ACSF: SR-95531 (1 µM, Tocris), CGP-55845 (1 µM, Tocris), kynurenic acid (2 mM, Sigma), and tetrodotoxin (TTX; 1 µM, Tocris) to block GABA_A_-receptor-, GABA_B_-receptor-, AMPA/NMDA-receptor- and sodium channel-mediated currents, respectively. CNO (5 µM; Sigma-Aldrich) was used to activate DREADDs. PGB (500 µM; Pfizer, New York, NY, USA) was used to examine its potential effects on PBN-SST neurons synaptic transmission.

### Behavioral tests

Mice were handled for at least 3 days before behavioral tests (Hurst and West, 2010). All behavioral tests were conducted during the light period of the light-dark cycle. Mice were moved to the behavioral room with dim light at least 30 min before experiments. In chemogenetic experiments, CNO was freshly dissolved in injection saline (10% v/v DMSO in 0.9% NaCl) and i.p. injected at 5 mg/kg of body weight. Behavioral tests were performed approximately 50 min after CNO injection.

#### Von Frey filament test

Mechanical hypersensitivity was assessed using the von Frey filament test. A series of von Frey filaments of increasing stiffness (0.04–1.4 g) were applied to the plantar surface of both hindpaws. Each filament was applied five times and the threshold (g) was taken as the lowest force that caused at least three withdrawals out of the five stimuli (Blackburn-Munro and Jensen, 2003; Hao et al., 1999).

#### Marble burying test

The clean cage (height, 12.5 cm; length, 28 cm; width, 13 cm) was filled with approximately 6 cm height of bedding. Twenty-four glass marbles (approximately 1.5 cm in diameter) were evenly spaced on top of the bedding with about 3 cm distance between each pair of marbles. Mice were placed individually into the cages and left undisturbed for 30 min (Chang et al., 2017). Marbles were considered buried if at least two-thirds of their surface was covered by bedding.

#### Light/dark box test

The test apparatus consisted of a two-compartment light-dark box (height, 42 cm; length, 42 cm; width, 42 cm) connected by a central opening at the floor level. Mice were placed individually in the center of the brightly lit side of the box and left undisturbed for 10 min of exploration. Transitions between the two compartments and total locomotor activity were recorded using the Tru-scan 2.0 system (Coulbourn Instrument, Allentown, PA, USA).

#### Elevated plus maze test (EPM)

The EPM is a common anxiety test that produces approach avoidance conflict (Walf and Frye, 2007). The EPM apparatus consisted of two open arms (length, 30 cm; width, 5 cm) and two closed arms (length, 30 cm; width, 5 cm; height, 25 cm) extending from the intersection zone (5 cm × 5 cm). The EPM was elevated 50 cm from the floor. The recording camera was placed above the maze. Mice were placed in the center of the intersection zone and then allowed to freely explore for 10 min. The open-arm time and total travel distance were measured with the video tracking software EthoVision XT 13 (Noldus Information Technology, Leesburg, VA, USA).

#### Sociability test

The three-chamber sociability test has been successfully employed to study social affiliation in several mouse lines (Yang et al., 2011). The social approach apparatus was an open-topped plastic box (height, 22 cm; length, 52.5 cm; width, 42.5 cm) divided into three chambers by two clear walls. The center compartment (length, 16.5 cm) was smaller than the other two compartments, which were equal in size to each other (length, 18 cm). The dividing walls had retractable doorways, allowing access into each chamber. A wire cup (bottom diameter, 5 cm) was used to contain the novel mice. Mice were housed alone for 24 h before the test and underwent the test in a darkened room. The lighting in the two side chambers was maintained at approximately 5 to 6 lux calibrated by a hand-held lux meter. Mice were habituated to the inverted wire cup for two 15 min sessions before the test session. Test mice were confined in the center chamber at the beginning of each phase for 10 min for habituation. During the habituation phase, each of the two side chambers contained an inverted empty wire cup. To initiate each 10 min phase, the doorways to the side chambers were opened, and the mice were allowed to explore freely. During the sociability phase, an unfamiliar mouse was enclosed in one of the wire cups in the side chambers. The time spent in each chamber and time spent exploring the enclosed novel mice or empty cups were recorded with a camera mounted overhead and analyzed using the EthoVision XT software 13 (Noldus Information Technology, Leesburg, VA, USA).

#### Forced swim test (FST)

The FST consisted of a transparent acrylic cylindrical container (height, 25 cm; width, 10 cm) filled with water to a height of 16 cm (Can et al., 2012; Castagné et al., 2010; Slattery and Cryan, 2012). The mouse was placed in the water maintained at 20–22 °C for 6 min. We measured the immobility time and frequency that were indicative of depression using the video tracking software EthoVision XT 13 (Noldus Information Technology, Leesburg, VA, USA).

### Data analysis and statistics

Data were analyzed using Clampfit 10.3 (Molecular Devices, Sunnyvale, CA, USA), custom-made program written in Python and GraphPad Prism 6 (GraphPad Software, Inc., San Diego, CA, USA). Statistical significance was tested using two-way ANOVA with Tukey’s *post hoc* test, two-way ANOVA with Bonferroni’s multiple comparisons test, Kolmogorov-Smirnov test, Wilcoxon matched-pairs signed rank test, or the Mann-Whitney test at the significance level (*p*) indicated. Data are presented as mean ± standard error of mean (SEM). Significance levels were set at *p* < 0.05 (*), *p* < 0.01 (**), *p* < 0.001 (***), or *p* < 0.0001 (****).

## Acknowledgments

We thank Drs. S.C. Lin, P.H. Chiang (National Yang Ming Chiao Tung University, Taiwan), A. Dominique (University of Liege, Belgium), and P. Turko (Charite University Medicine Berlin, Germany) for commenting on earlier versions of the manuscript and all the members of the Lien lab for insightful discussions. We also thank Pfizer for supply of pregabalin.

## Funding

This work was financially supported by the Brain Research Center, National Yang Ming Chiao Tung University from the Featured Areas Research Center Program within the framework of the Higher Education Sprout Project by the Ministry of Education in Taiwan, National Health Research Institutes (NHRI-EX109-10814NI), and Ministry of Science and Technology (MOST 108-2321-B-010-009-MY2, MOST 108-2320-B-010-026-MY3, MOST 108-2638-B-010-002-MY2, MOST 109-2926-I-010-506, MOST 110-2321-B-010-006, MOST 111-2321-B-A49-005) in Taiwan.

## Author contributions

Y.L.L., W.Y.W., Z.S.Y., and C.C.L., conceptualized the project and wrote the paper. Y.L.L., W.Y.W., Z.S.Y., executed the experiments. Y.L.L., W.Y.W., Z.S.Y., S.C.L., analyzed the data. J.K.C., S.P.C., H.L. and S.J.W., conceptualized the project and interpreted the data, and C.C.L., acquired funding.

## Competing Interests

The authors declare that pregabalin was provided from Pfizer. There are no potential conflicts of interest with respect to the research, authorship, and publication of this article.

## Data Availability

Source data file contains the numerical data used to generate the figures and perform the statistics.

---

**Figure 1 - Figure supplement 1.**
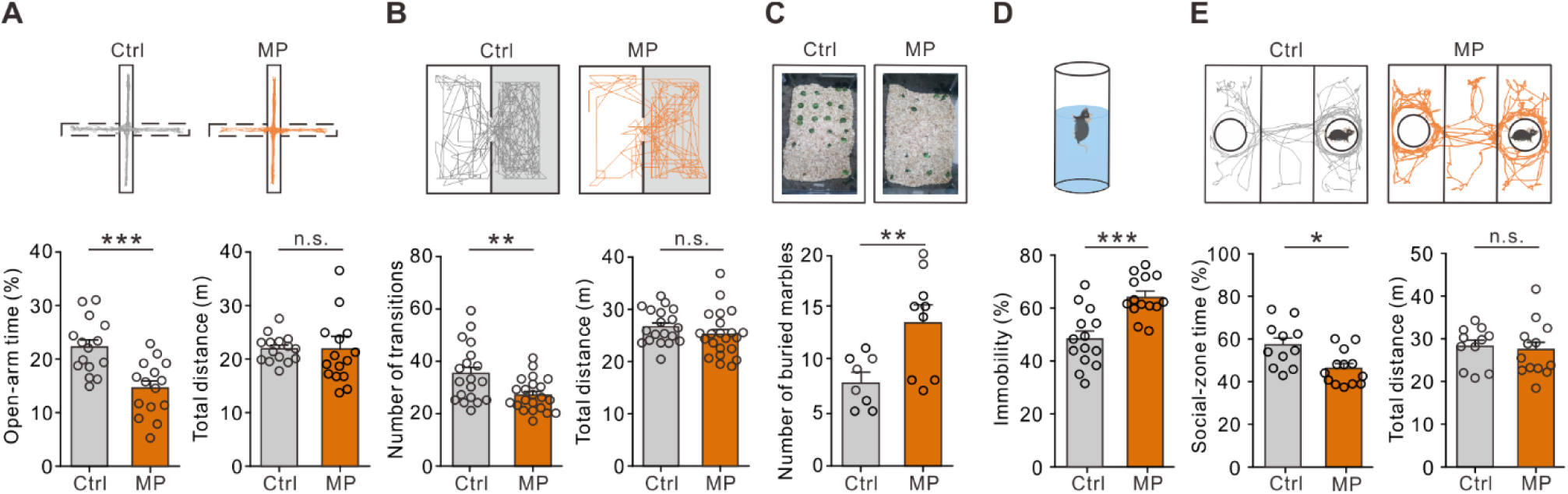
Comorbid affective symptoms in a mouse model of MP. **(A)** Top, representative travel trajectories of each group during the EPM test. Bottom, summary plots of open-arm time (Ctrl, 22.3 ± 1.3%, n = 15; MP, 14.7 ± 1.3%, n = 15; Mann-Whitney test, U = 33, \*\*\**p* = 0.001) and total distance (Ctrl, 21.9 ± 0.6 m, n = 15; MP, 21.9 ± 1.6 m, n = 15; Mann-Whitney test, U = 95, n.s., non-significant, *p* = 0.495). **(B)** Top, representative travel trajectories of each group during the L/D box test. Bottom, number of transitions (Ctrl, 35.5 ± 2.4, n = 19; MP, 26.6 ± 1.5, n = 22; Mann-Whitney test, U = 104.5, \*\**p* = 0.005) and total distance (Ctrl, 26.1 ± 1.6 m, n = 19; MP, 25.3 ± 1.0 m, n = 22; Mann-Whitney test, U = 169, n.s., non-significant, *p* = 0.303). **(C)** Top, representative results of the marble burying test. Bottom, summary plot of the number of buried marbles (Ctrl, 7.9 ± 0.9, n = 8; MP, 13.6 ± 1.6, n = 9; Mann-Whitney test, U = 12.5, \*\**p* = 0.018). **(D)** Top, schematic of the FST. Bottom, summary plot of immobility time (Ctrl, 48.7 ± 2.9%, n = 14; MP, 64.7 ± 2.1%, n = 14; Mann-Whitney test, U = 23, \*\*\**p* = 0.0003). **(E)** Top, representative trajectories of each group during the three-chamber sociability test. Bottom, summary plots of relative time in the social zone (Ctrl, 57.3 ± 3.2%, n = 11; MP, 46.3 ± 2.2%, n = 13; Mann-Whitney test, U = 28, \*\**p* = 0.011) and total distance (Ctrl, 28.2 ± 1.4 m, n = 11; MP, 27.4 ± 1.8 m, n = 13; Mann-Whitney test, U = 66, n.s., non-significant, *p* = 0.766).

**Figure 3 - Figure supplement 1.**
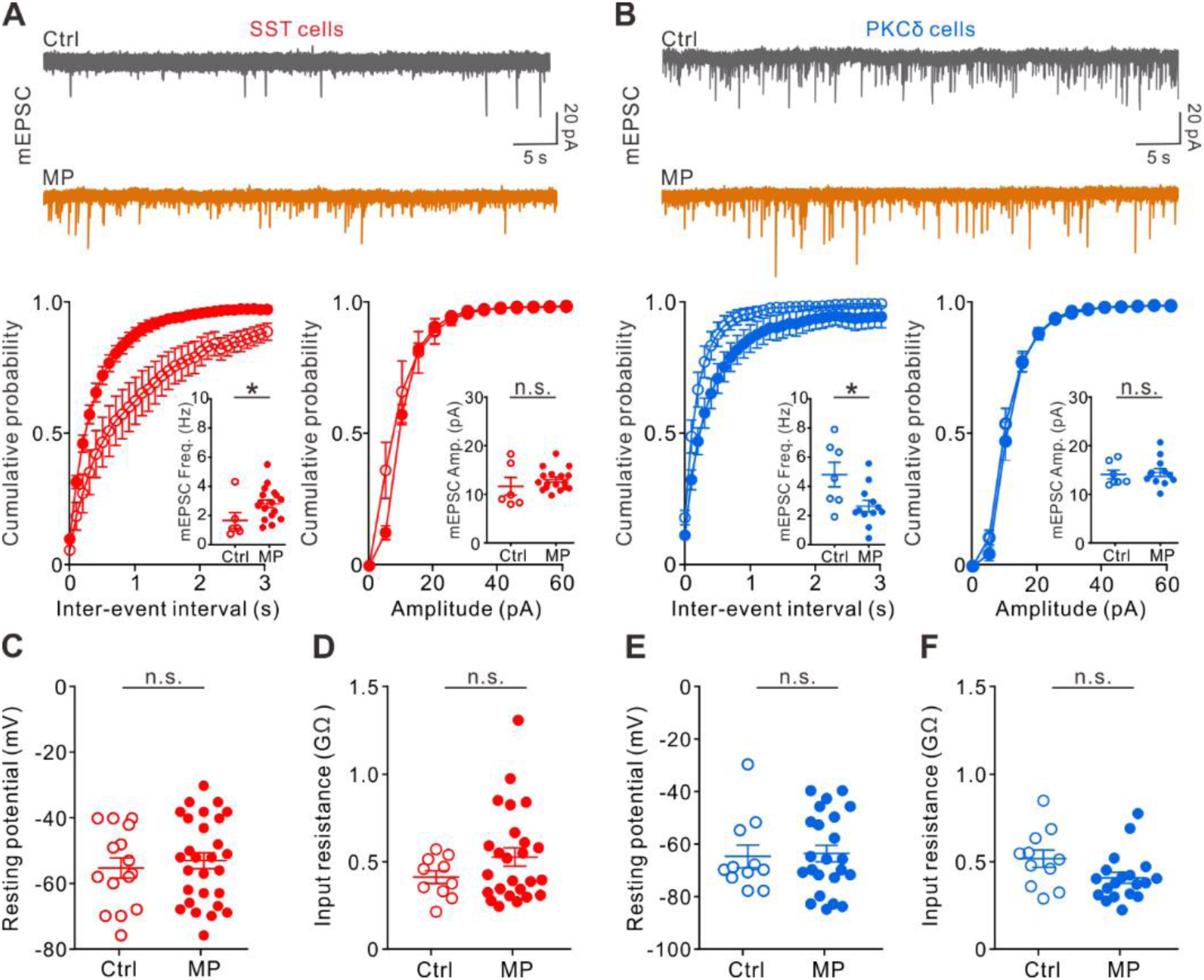
Comparison of mEPSC, resting potential, and input resistance of CeA neurons in Ctrl and MP mice. **(A)** Top, representative traces of mEPSCs recorded from CeA-SST neurons of Ctrl and MP mice. Bottom, cumulative probability of inter-event interval (Ctrl, n = 6; MP, n = 17; Kolmogorov-Smirnov test, \*\*\**p* < 0.0001. Inset, summary of mEPSC frequency, Ctrl, 1.4 ± 0.5 Hz, n = 6; MP, 2.5 ± 0.3 Hz, n = 17, Mann-Whitney test, U = 20, \**p* = 0.029) and amplitude (Ctrl, n = 6; MP, n = 17; Kolmogorov-Smirnov test, n.s., non-significant, *p* = 0.518. Inset, summary of mEPSC amplitude, Ctrl, 11.8 ± 1.8 pA, n = 6; MP, 13.2 ± 0.5 pA, n = 17, Mann-Whitney test, U = 33, n.s., non-significant, *p* = 0.215). **(B)** Top, representative traces of mEPSCs recorded from CeA-PKCδ neurons of Ctrl and MP mice. Bottom, cumulative probability of inter-event interval (Ctrl, n = 7; MP, n = 12; Kolmogorov-Smirnov test, \*\*\**p* = 0.0002. Inset, summary of mEPSC frequency, Ctrl, 4.8 ± 0.8 Hz, n = 7; MP, 2.6 ± 0.4 Hz, n = 12, Mann-Whitney test, U = 17, \**p* = 0.036) and amplitude (Ctrl, n = 7; MP, n = 12; Kolmogorov-Smirnov test, n.s., non-significant, *p* = 0.993. Inset, summary of mEPSC amplitude, Ctrl, 14.0 ± 0.9 pA, n = 7; MP, 14.6 ± 0.8 pA, n = 12, Mann-Whitney test, U = 34, n.s., non-significant, *p* = 0.514). **(C)** Summary of resting potential of CeA-SST neurons (Ctrl, -55.3 ± 3.2 mV, n = 15; MP, -53.0 ± 2.5 mV, n = 28; Mann-Whitney test, U = 181, n.s., non-significant, *p* = 0.468). **(D)** Summary of input resistance of CeA-SST neurons (Ctrl, 0.42 ± 0.04 GΩ, n = 10; MP, 0.53 ± 0.05 GΩ, n = 25; Mann-Whitney test, U = 97, n.s., non-significant, *p* = 0.312). **(E)** Summary of resting potential of CeA-PKCδ neurons (Ctrl, -64.9 ± 4.3 mV, n = 11; MP, -63.8 ± 3.1 mV, n = 23; Mann-Whitney test, U = 120, n.s., non-significant, *p* = 0.821). **(F)** Summary of input resistance of CeA-PKCδ neurons (Ctrl, 0.52 ± 0.05 GΩ, n = 11; MP, 0.41 ± 0.03 GΩ, n = 19; Mann-Whitney test, U = 60, n.s., non-significant, *p* = 0.057).

**Figure 4 - Figure supplement 1.**
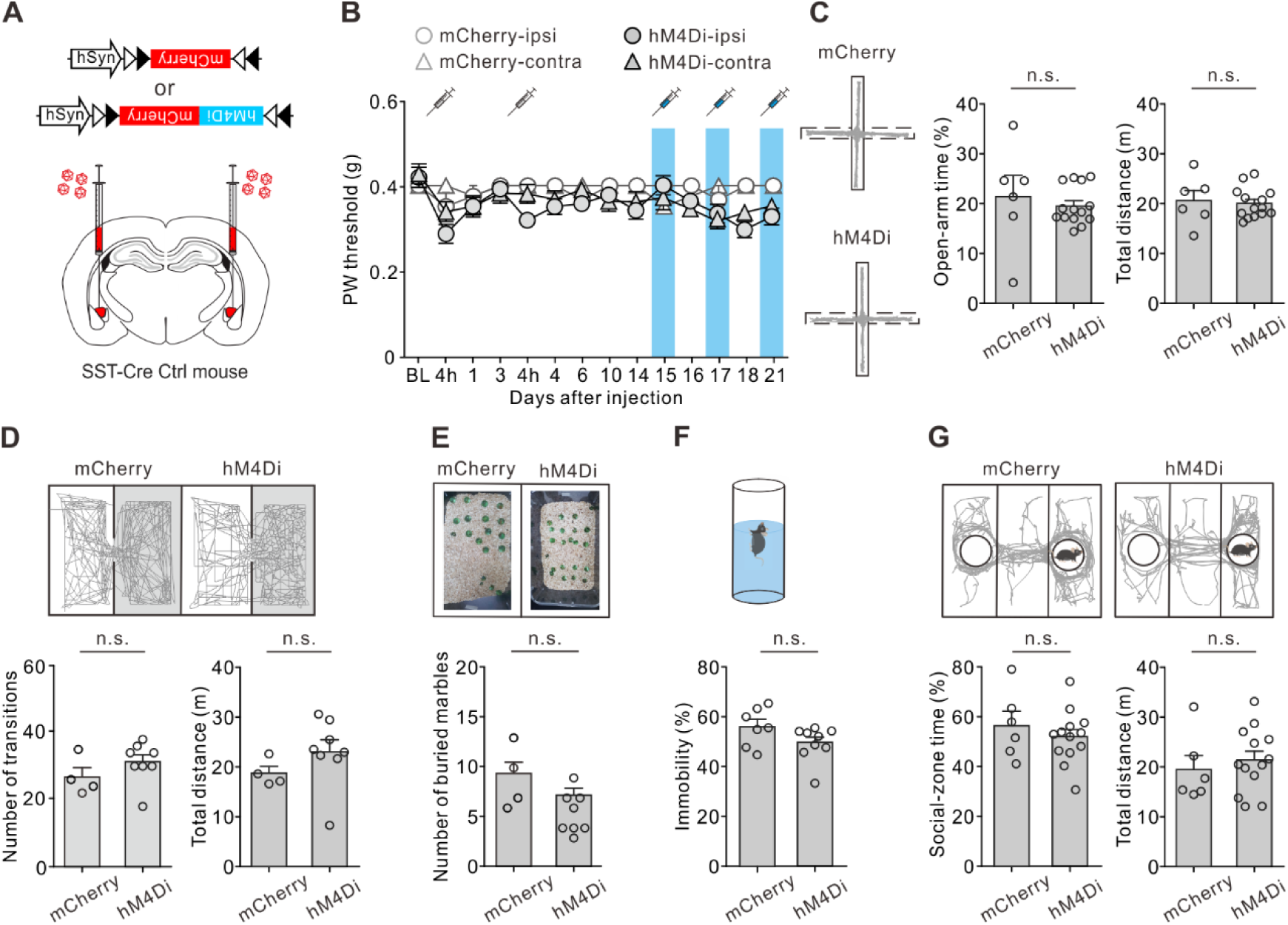
Suppressing CeA-SST neuron excitability in Ctrl mice exerted little effect on nociception and affective behaviors. **(A)** Viral constructs and experimental schematic. **(B)** Effect of CNO treatment on PW threshold in Ctrl mice (mCherry, n = 10; hM4Di, n = 52; two-way ANOVA with Tukey’s post hoc test, F(3,1616) = 1.22, *p* = 0.26). Blue area indicates the period of CNO treatment. **(C)** Left, representative trajectories during the EPM test. Right, summary of the effects of CNO treatment on the open-arm time (mCherry, 21.3 ± 4.2%, n = 6; hM4Di, 19.5 ± 1.0%, n = 14; Mann-Whitney test, U = 34, n.s., non-significant, *p* = 0.545) and total distance (mCherry, 20.7 ± 2.0 m, n = 6; hM4Di, 20.1 ± 0.8 m, n = 14; Mann-Whitney test, U = 37, n.s., non-significant, *p* = 0.716). **(D)** Top, representative travel paths of each Ctrl group during the L/D box test. Bottom, summary of CNO effect on the number of transitions (mCherry, 26.8 ± 2.9, n = 4; hM4Di, 31.3 ± 2.2, n = 8; Mann-Whitney test, U = 9, n.s., non-significant, *p* = 0.259) and total distance (mCherry, 18.7 ± 1.4 m, n = 4; hM4Di, 23.0 ± 2.4 m, n = 8; Mann-Whitney test, U = 6, n.s., non-significant, *p* = 0.105). **(E)** Top, representative images of the marble burying test. Bottom, summary of the effects of CNO treatment on the number of buried marbles (mCherry, 7.8 ± 2.5, n = 4; hM4Di, 5.6 ± 0.8, n = 8; Mann-Whitney test, U = 5.5, n.s., non-significant, *p* = 0.083). **(F)** Top, schematic of the FST setup. Bottom, summary of relative time of immobility (mCherry, 56.1 ± 2.9%, n = 7; hM4Di, 49.2 ± 2.3%, n = 9; Mann-Whitney test, U = 15, n.s., non-significant, *p* = 0.091). **(G)** Top, representative travel paths during the three-chamber sociability test. Bottom, summary of the effects of CNO treatment on the social-zone time (mCherry, 56.6 ± 5.6%, n = 6; hM4Di, 52.0 ± 2.9%, n = 13; Mann-Whitney test, U = 32, n.s., non-significant, *p* = 0.577) and total distance (mCherry, 19.5 ± 2.7 m, n = 6; hM4Di, 21.3 ± 1.8 m, n = 13; Mann-Whitney test, U = 32, n.s., non-significant, *p* = 0.549).

**Figure 4 - Figure supplement 2.**
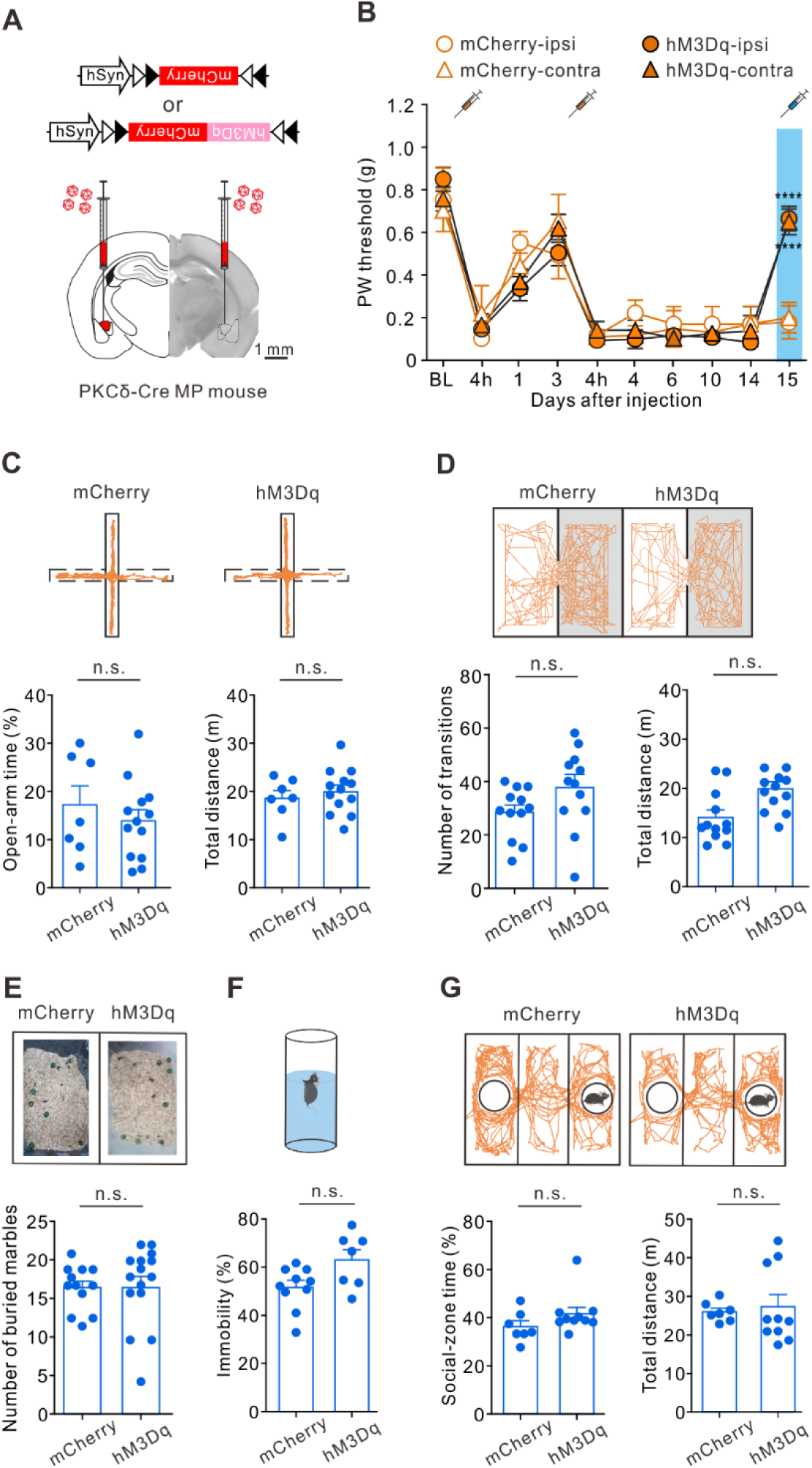
Enhancing PKCδ neuron excitability reduced pain, but failed to alleviate affective symptoms. **(A)** Viral constructs and experimental schematic. Scale bar, 1 mm. **(B)** Effect of CNO on PW threshold in hM3Dq-MP mice (mCherry, n = 4; hM3Dq, n = 13; two-way ANOVA with Tukey’s post hoc test, F(9,320) = 47.72, \*\*\*\**p* < 0.0001 relative to day 14. The blue area indicates the period of CNO treatment). **(C)** Top, representative trajectories of each MP group during the EPM test. Bottom, summary of the effects of CNO treatment on open-arm time (mCherry, 17.2 ± 3.9%, n = 7; hM3Dq, 13.9 ± 2.3%, n = 13; Mann-Whitney test, U = 39, n.s., non-significant, *p* = 0.639) and total distance (mCherry, 18.8 ± 1.6 m, n = 7; hM3Dq, 20.2 ± 1.3 m, n = 13; Mann-Whitney test, U = 39, n.s., non-significant, *p* = 0.616). **(D)** Top, representative travel paths of each MP group during the L/D box test. Bottom, summary of CNO effect on the number of transitions (mCherry, 28.3 ± 2.8, n = 12; hM3Dq, 37.1 ± 4.4, n = 12; Mann-Whitney test, U = 38.5, n.s., non-significant, *p* = 0.053) and total distance (mCherry, 16.0 ± 1.8 m, n = 12; hM3Dq, 14.0 ± 1.5 m, n = 12; Mann-Whitney test, U = 51, n.s., non-significant, *p* = 0.239). **(E)** Top, representative images of the marble burying test. Bottom, summary of the effects of CNO treatment on the number of buried marbles (mCherry, 15.8 ± 0.8, n = 12; hM3Dq, 16.1 ± 1.2, n = 15; Mann-Whitney test, U = 74.5, n.s., non-significant, *p* = 0.461). **(F)** Top, schematic of the FST setup. Bottom, summary of relative time of immobility (mCherry, 51.6 ± 2.8%, n = 10; hM3Dq, 63.3 ± 4.2%, n = 7; Mann-Whitney test, U = 16.5, n.s., non-significant, *p* = 0.074). **(G)** Top, representative travel paths of each MP group during the three-chamber sociability test. Bottom, summary of the effects of CNO treatment on social-zone time (mCherry, 36.1 ± 2.4%, n = 7; hM3Dq, 41.6 ± 2.6%, n = 10; Mann-Whitney test, U = 18, n.s., non-significant, *p* = 0.107) and total distance (mCherry, 26.0 ± 1.0 m, n = 7; hM3Dq, 27.3 ± 3.1 m, n = 10; Mann-Whitney test, U = 26, n.s., non-significant, *p* = 0.414).

**Figure 4 - Figure supplement 3.**
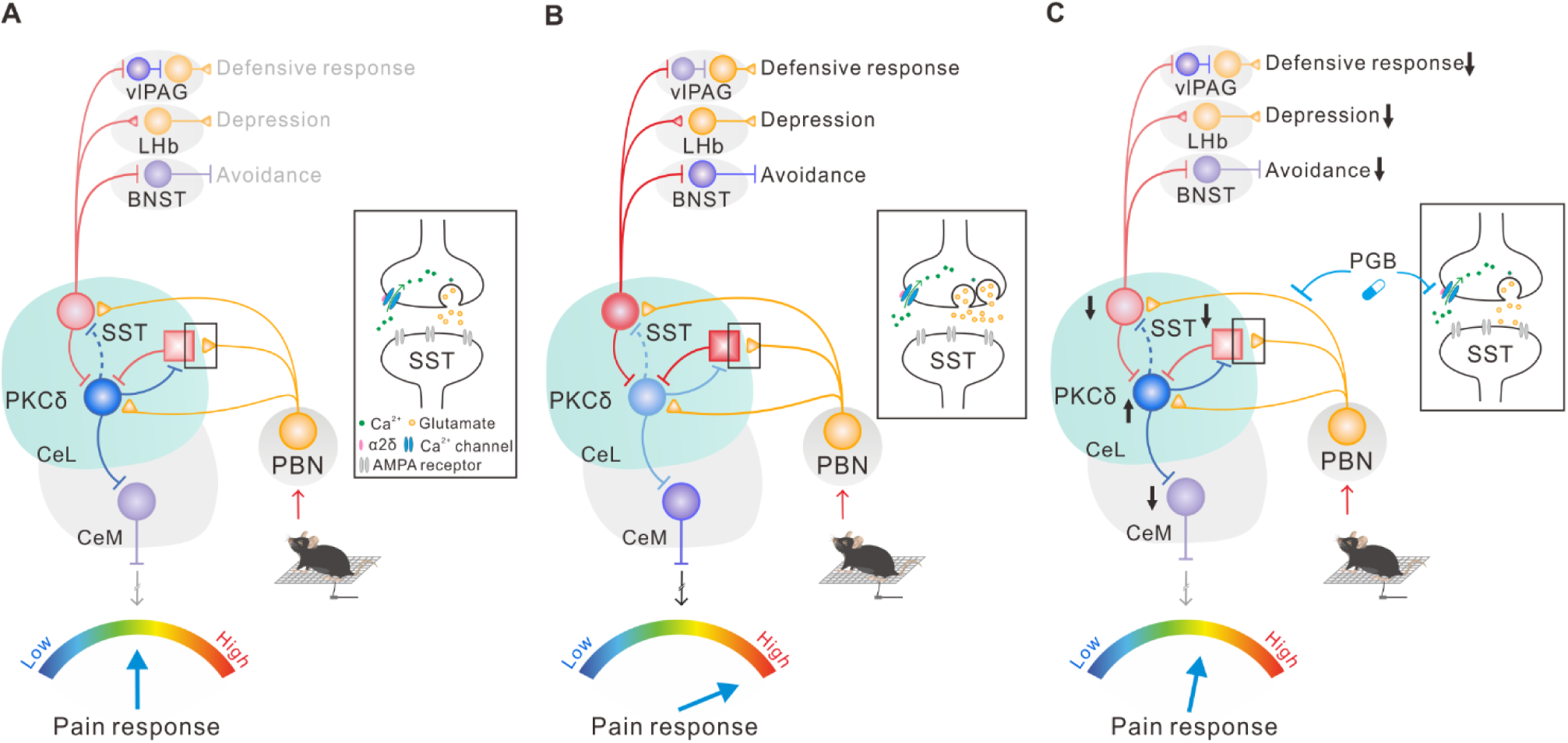
Proposed wiring diagram of CeA circuits, extrinsic pathways and behavioral responses. **(A-C)** CeA-SST neurons comprise at least two subpopulations. One subpopulation of CeA-SST neurons (red circle) project to the BNST for approach-avoidance behaviors, the midbrain vlPAG for active avoidance behavior, and the LHb for depression-like behavior. CeA-PKCδ neurons (blue circle) form reciprocal inhibition with another subpopulation of CeA-SST neurons (red square) and project to the major output of CeA, which is the CeM region, and then project to the PAG region for pain behavior. Both CeA-SST neuron subpopulations receive input from the PBN and target CeA-PKCδ neurons. **(A)** normal condition; **(B)** MP state; **(C)** treated state after chemogenetic inactivation of CeA-SST neurons or activation of CeA-PKCδ neurons or after PGB treatment.

## Notes

### Competing Interest Statement

The authors have declared no competing interest.

